# Pearling Drives Mitochondrial DNA Nucleoid Distribution

**DOI:** 10.1101/2024.12.21.629917

**Authors:** Juan C. Landoni, Matthew D. Lycas, Josefa Macuada, Willi Stepp, Roméo Jaccard, Christopher J. Obara, Andrew S. Moore, David Hoffman, Jennifer Lippincott-Schwartz, Wallace Marshall, Gabriel Sturm, Suliana Manley

## Abstract

The mitochondria of most eukaryotes carry an indispensable second genome (mtDNA), encoding genes engaged in oxidative phosphorylation^1^. The regular positioning and segregation of mtDNA-containing nucleoids is essential for mitochondrial function and inheritance, as well as cellular health^2–5^. However, the underlying mechanism driving nucleoid distribution and disaggregation remains unknown^6,7^. Our data reveal that mitochondria frequently undergo reversible pearling, a biophysical instability that undulates tubules into regularly spaced beads ^8^, typically triggered by calcium influx. We discovered that physiological pearling imposes a characteristic length scale, simultaneously mediating nucleoid disaggregation and establishing inter-nucleoid distancing with near-maximally achievable precision. We found that lamellar cristae invaginations of the inner mitochondrial membrane play a dual role, determining pearling frequency and duration, and preserving the resulting nucleoid spacing after organelle recovery to a tubular form. Thus, disrupting cristae ultrastructure resulted in more frequent pearling, but also aberrant nucleoid clustering. Our results demonstrate that the distribution of mitochondrial genomes is governed by the interplay between rapid and reversible pearling and cristae ultrastructure, establishing a mechanism for this long-puzzling yet fundamental feature of eukaryotic life, and offering insights into its potential modulation.

## Introduction

Mitochondria carry their own independently replicating genome, mitochondrial DNA (mtDNA), which is considered a remnant of their endosymbiotic origins^9^. In humans, mtDNA encodes 13 core subunits of the oxidative phosphorylation system, as well as RNAs required for their expression. Consequently, mtDNA integrity and copy number maintenance are critical for mitochondrial function and cellular health, with their disruption leading to a range of metabolic and neurological disorders^1,10^. Each human cell contains hundreds to thousands of mtDNA copies, packaged into nucleoids and locally confined by inner membrane cristae invaginations^11,12^.

Regular spacing enables precise inheritance of mtDNA by daughter cells during division^2,13,14^, and permits the well-distributed gene expression along the mitochondrion^3,15^. However, no molecular machinery has thus far been identified to segregate nucleoids or maintain their regular distancing^6^. Defects in mitochondrial dynamics can disrupt nucleoid homeostasis^7,16^, but mitochondria can still maintain nucleoid distribution in the absence of fission and fusion and fission and tethering in yeast and humans respectively^17–19^, indicating that the underlying mechanism remains to be discovered.

Our data indicate that the mitochondrial *pearling* instability —a shape transition from a tubule into beads-on-a-string— is a rapid, frequent, and typically reversible event in eukaryotic mitochondrial dynamics. Reminiscent of the Plateau-Rayleigh instability driven by competition between elasticity and surface tension^8,20^, these simultaneously occurring, regularly spaced constrictions have been reported anecdotally^21–23^ and described as priming steps for fission^24,25^. Here, we combine super-resolution microscopy approaches and genetic modulation of cristae morphology, membrane tethering, fission, and calcium dynamics to demonstrate the emergent functional role of *pearling* in redistributing mtDNA copies and establishing the long enigmatic regular distancing of nucleoids.

### Mitochondrial pearling is a physiological and reversible biophysical instability

Mitochondrial pearling has typically been observed following its induction (*i*.*e*. by membrane rupture, calcium ionophore treatment, or mitochondrial hyper-elongation^20–25^). To assess its relevance for homeostatic processes, we sought to observe its occurrence in steady-state conditions. Spontaneous pearling events, defined as those arising in a cell without physical or pharmacological induction, occurred in untreated human U2-OS cells (Fig 1a, Supplementary Movie 1) with an incidence of ~1.5 ± 0.4 events/cell/min under widefield fluorescence imaging. Reversible pearling emerged rapidly from one frame to the next (when imaging in phase contrast up to 100 frames-per-second) and lasted 5-30 seconds, with no apparent subcellular location bias (Extended Data Fig. 1a). Consistent with previous reports^22^, pearling often ended with a fission at one or more of the constrictions (Supplementary Movie 1). The mean pearl diameter as measured by a matrix label was 425 ± 86 nm (Fig. 1b, Extended Data Fig. 1b), and the median number of pearls per event was five, although we observed strings of up to 34 pearls (Extended Data Fig. 1c). Reversible mitochondrial pearling was conserved across a range of mammalian cell types and the yeast *Saccharomyces cerevisiae* (Extended Data Fig. 1c) with comparable morphologies and dynamics^26^, and across an array of labelling strategies (outer mitochondrial membrane (OMM)- and matrix-targeted fluorescent proteins, vital stains for matrix and the inner mitochondrial membrane (IMM), and label-free contrast), and microscopy techniques (widefield epi-fluorescence, light sheet, instant structured illumination (iSIM), stimulated emission depletion (STED)) (Fig. 1a-e & Extended Data Fig. 1c). Additionally, we captured pearling using phase contrast imaging, both spontaneously and upon ionomycin induction (Extended Data Fig. 1c-d), as well as using an event-driven acquisition paradigm (modified from^27^). Briefly, a neural network trained to predict pearling events in phase contrast triggered real-time fluorescent imaging only when pearling was detected, minimizing phototoxicity or potential laser-related induction (Extended Data Fig. 1e). Taken together, our data indicates that pearling is a physiological, frequent and evolutionarily conserved shape transformation in mitochondrial dynamics, constricting OMM, IMM, and matrix (Fig. 1f).

**Figure 1.**
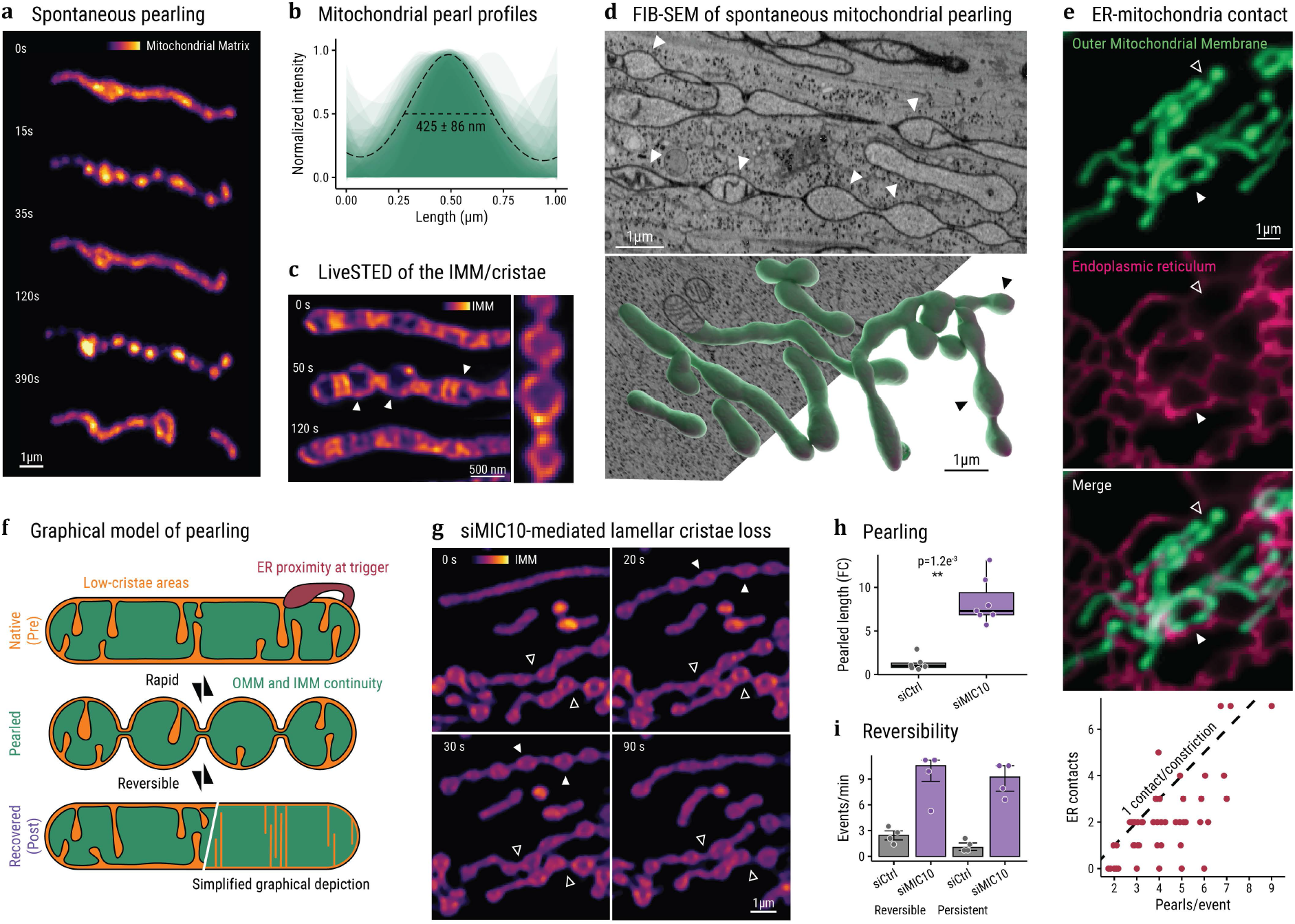
Spontaneous mitochondrial pearling. **(a)** U2-OS cell mitochondrion (mitochondria-targeted mt-StayGold) undergoing consecutive reversible pearling (15 s, 120 s), and fission (390 s), iSIM. **(b)** Line profiles of 50 spontaneous pearls (as in (a)), with the average full-width half-max diameter. **(c)** Time-lapse and snapshot of PKMO-labeled IMM cristae during spontaneous pearling (arrowheads) in live STED microscopy (n = 57 timelapses over 4 independent experiments). **(d)** FIB-SEM image and 3D reconstructions of mitochondria (arrowheads select spontaneous pearls in an uninduced U2-OS cell). **(e)** OMM (TOM20-mScarlet, green) and endoplasmic reticulum (ER-StayGold, pink), highlighting pearling constrictions with (filled arrowhead) or without (empty arrowhead) mitochondria-ER proximity. Below, quantified number of ERMCS versus the number of pearls. Each jittered point represents an independent pearling event; the dashed line represents one ERMCS at every pearl constriction. **(f)** Graphical model of mitochondrial pearling. **(g)** siRNA-mediated MIC10 silenced HeLa cells, highlighting rapidly reversible (filled arrowheads) or persistent (empty arrowheads) pearling events. Array-detector confocal imaging of PKMO-stained mitochondria (n = 86 control + 91 siMIC10 cells, from 4 independent experiments). **(h)** Pearling in siRNA control (n = 6) and siMIC10 (n = 7) cells as in (g): proportion of the total mitochondrial network length which pearls within 1 minute, fold-change to median of control cells. **(i)** Mean reversible pearling event rate per minute per experiment (n = 4 independent experiments), defined by pearling completed within a minute or the end of acquisition time. Bar and box plots represent median ± IQR of the overlaid data points, p values from two-sided Wilcoxon test. OMM = outer mitochondrial membrane, IMM = inner mitochondrial membrane, siMIC10 = siRNA-mediated MIC10 silencing, siCtrl = scrambled siRNA control.

To investigate the finer ultrastructure of pearling, we turned to electron microscopy of U2-OS cells preserved in a near-native state by high-pressure freezing. Following freeze-substitution and subsequent focused ion beam-scanning electron microscopy (FIB-SEM) (Fig. 1d, Extended Data Fig. 1f), we observed that both IMM and OMM remain continuous during pearling^25^, leaving the matrix connected but tightly constricted. Our data presented a relatively low density of cristae, especially in pearled mitochondria (Fig. 1d), compared with other reports in HeLa cells^28^. Live STED imaging of the IMM corroborated this observation, with lower cristae density in U2-OS cells than HeLa cells (Extended Data Fig. 1g), and pearls frequently devoid of cristae signal (Fig. 1c). We observed that ~80% of pearling events began with contact between a mitochondrion and an ER tubule somewhere along the mitochondrion, but many constrictions of the pearled mitochondrion lacked a contact (Fig. 1e, ^26^). This indicates the role of ER-mitochondria contact sites (ERMCS) to be in triggering pearling through signaling, consistent with previous reports^24,25^, rather than physically stabilizing constrictions.

Physically, a membrane tube remains stable as long as its elastic resistance to deformation dominates over its membrane tension, which otherwise drives spherical shapes^8,20^. The elasticity of a mitochondrion arises from its membranes and cristae folds^20,29^, so we expected that modulating cristae density could modify pearling onset and stability, as suggested by our IMM and EM imaging (Fig. 1c-d). While most cristae perturbations severely disrupt mitochondrial homeostasis and lead to fragmentation, removal of the mitochondrial contact site and cristae organizing system (MICOS) core subunit MIC10 in HeLa cells depletes lamellar cristae with mild-to-negligible effects on respiratory chain assembly or network morphology^30^. Depletion of MIC10 (siMIC10) in cristae-dense HeLa cells resulted in a striking increase in spontaneous pearling probability when compared to siRNA controls, as measured by increases in the proportion of mitochondria pearled per minute (Fig. 1g-h, Extended Data Fig. 1h), the number of individual events per minute (Fig. 1i), and the percentage of cells with pearled mitochondria in a single snapshot (from 17 ± 4 % to 50 ± 11%). In addition to frequent reversible events, persistent pearling events lasting longer than a minute or even beyond the entire 6-minute acquisition were abundant in siMIC10 cells (Fig. 1i). This supports our model that lamellar cristae elastically resist the pearling instability, both diminishing its prevalence and enabling rapid recovery to a tubular organelle form.

### Pearling entrains mtDNA nucleoids

As is characteristic of Rayleigh-Plateau-like instabilities, mitochondrial pearling presents undulations of a characteristic wavelength, in this case a mean distance between pearls of 0.81±0.13 μm as measured by STED microscopy (Fig. 2a). This distance is nearly identical to the median spacing of mtDNA nucleoids we measure along 3D mitochondrial networks in U2-OS cells with light sheet microscopy (0.84 ± 0.13 μm, Fig. 2b) and published values in yeast^2,18^, which prompted us to investigate the relationship between pearling and nucleoids.

**Figure 2.**
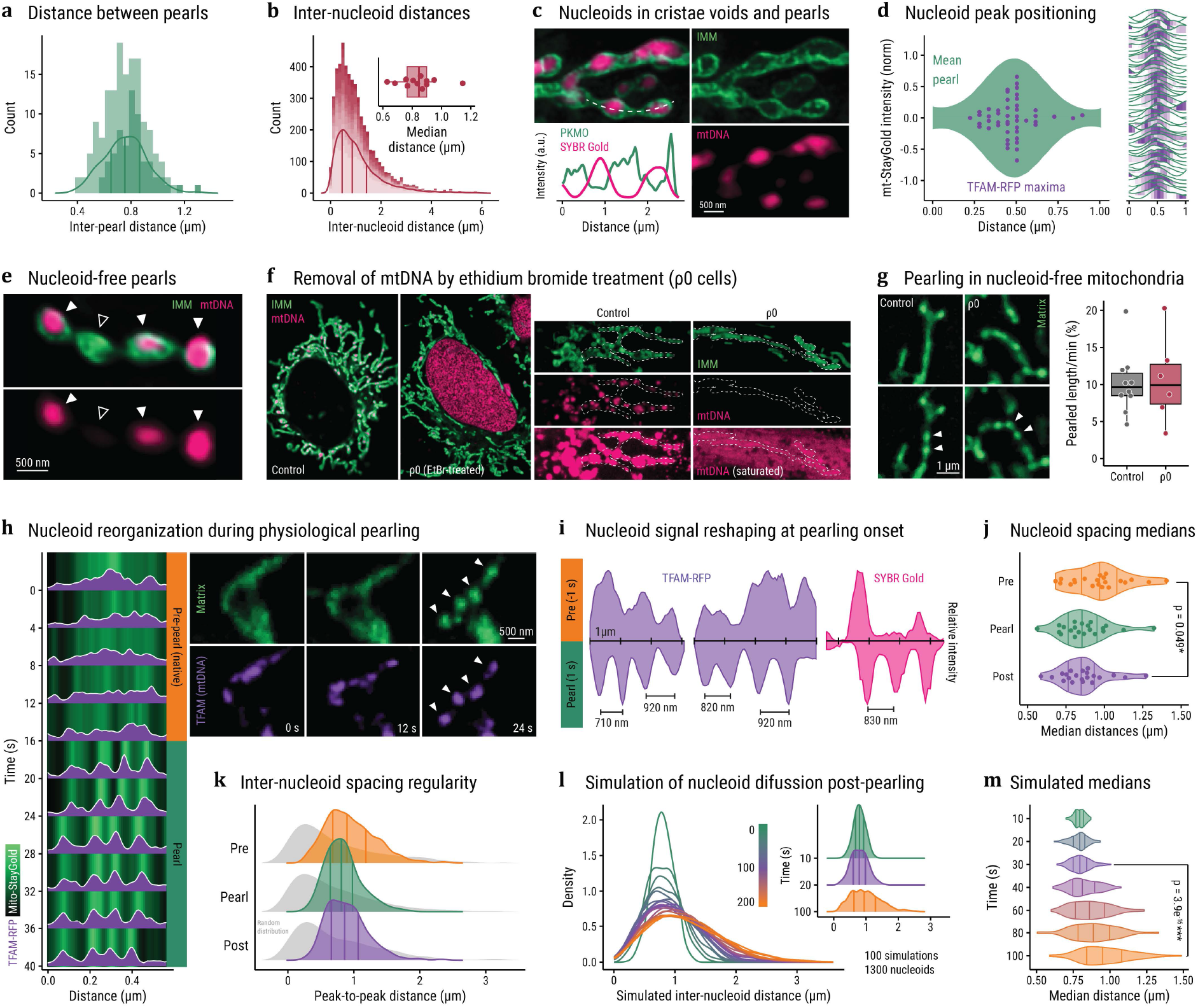
Pearling-driven mtDNA redistribution. **(a)** Distribution of inter-pearl distances, as measured in U2-OS cells by iSIM and STED (dim and dark histograms respectively), overlaid with a density curve and vertical median and inter-quartile range lines (n = 87+63 inter-pearl distances). **(b)** Inter-nucleoid distance distribution along 3D light sheet images of in U2-OS mitochondria and nucleoids (PKMO and SYBR gold staining). Different cells are displayed as different shades in the stacked histogram, each cell’s medians as inset (n = 5319 inter-nucleoid distances in 12 cells). **(c)** Live imaging of IMM (STED, PKMO, green) and nucleoid (Confocal, SYBR gold, pink). Line profile (bottom left) displaying relative intensities along the dashed line (U2-OS cells, representative from n = 4 independent experiments). **(d)** Position of nucleoid maxima (TFAM-RFP, indigo) displayed relative to the average profile of a pearl (mt-StayGold, mirrored). On the right, individual pearl profiles with nucleoid intensity heatmap in indigo (n = 50 pearls). **(e)** Spontaneous pearling as in (c), with nucleoid-occupied (filled arrowheads) and a nucleoid-free pearl (empty arrowhead). **(f)** Array-detector confocal imaging of IMM (PKMO, green) and DNA (SYBR gold, pink) in control and ethidium bromide-treated conditions (mtDNA-free mitochondria). SYBR gold staining of the nucleus in ρ0 cells is an expected consequence of mtDNA depletion. **(g)** Spontaneous pearling captured in control and ρ0 cells, and pearling frequency quantified as the percentage of the total mitochondrial network length pearling per minute. n = 2 independent experiments. **(h)** Timelapse of spontaneous pearling as in (d), showing nucleoid (TFAM-RFP, indigo) signal condensation concomitant with mitochondrial (mt-StayGold, green) pearling onset, and corresponding kymograph and line profiles. **(i)** Normalized mtDNA intensity line profile examples, from TFAM-RFP (indigo) or SYBR gold (pink) signal immediately before and after pearling onset (mirrored). **(j-k)** Distribution of inter-nucleoid distances during pearling (pearl) and within 10s before (pre) and after (post) pearling onset (k), including simulated random distributions for each population (gray density curves), and median distance values per independent pearling event (j). Wilcoxon test, n = 21 independent events with an average of 5.5 nucleoids and 15 time points each. **(l-m)** Distance distributions obtained from 100 post-pearling nucleoid diffusion simulations. Each line represents a measured timepoint between 0 and 200 s (l), inset with highlighted distributions colored to match their experimental distribution in (k), and median inter-nucleoid distances per simulated event (m). Lines in density and violin plots represent the median and IQR of the distribution, Wilcoxon test. IMM = inner mitochondrial membrane, mtDNA = mitochondrial DNA, ρ0 = Ethidium bromide-treated mtDNA-free.

Dual-color imaging of nucleoids and IMM by STED or matrix by iSIM revealed that most pearls contained a nucleoid, located near their center (Fig. 2c-d, Supplementary Movie 2). However, pearls spontaneously occurred without nucleoids (Fig. 2e, Extended Data Fig. 2a), and acute depletion of mtDNA by ethidium bromide (Extended Data Fig. 2b) led to the disappearance of nucleoids but did not affect network and cristae morphology (Fig. 2f) or pearling frequency or form (Fig. 2g). Thus, nucleoids do not determine pearling onset, shape or positioning.

Instead, we speculated that the converse may be true – that pearling may govern nucleoid positioning. Using live iSIM, we assessed nucleoid organization before, during, and after mitochondrial pearling events under non-induced conditions. In contrast to their sometimes diffuse or irregular organization pre-pearling, nucleoids within pearls appeared as distinct puncta (Fig. 2h-i) with regular spacing consistent with population averages. We replicated these results using genetically encoded (mt-StayGold and TFAM-RFP) and dye-based probes (SYBR gold and MitoTracker Red FM), excluding possible effects of dye intercalation or protein overexpression. The onset of pearling can reshape and redistribute nucleoids, resulting in more discrete and regularly spaced foci.

To collect statistics on pearling-driven nucleoid redistribution, we measured the distances between neighboring nucleoids before, during, and after pearling in a larger dataset from widefield microscopy. Pearling-associated nucleoid spacing replicated the non-random spacing from light sheet data (Fig. 2b) and previous reports^2,18^, peaking at 800-900 nm (Fig. 2j-k). Moreover, the onset of pearling leads to a narrower and more symmetrical distribution when compared to pre-pearling. Importantly, post-recovery to a native tubular shape, the distribution remains narrower (Fig. 2k). The median distances between nucleoids per event are significantly smaller (Fig. 2j) during and after pearling compared to before. While this reduction may seem counter-intuitive, it is likely due to the disaggregation of larger nucleoids, as explored subsequently.

To test our model, we generated a simple diffusion simulation for mtDNA nucleoids post-pearling. We reasoned that immediately following a pearling event, nucleoids take on post-pearling experimental distances, and then diffuse until the next pearling event to take on pre-pearling distances. In short, we initialized nucleoids spaced equidistantly by 810 nm within a tubular container to represent a mitochondrion immediately following pearling recovery (400 nm x 10 μm, (Extended Data Fig. 2c-d). We simulated 2D diffusion using published instantaneous nucleoid velocities^31^ and, to account for imaging resolution and nucleoid aggregation, we merged nucleoids within 120 nm of one another. Following 100 simulations, the inter-nucleoid distance distributions as well as the medians per simulation closely replicate those experimentally observed (Fig. 2l-m), with experimental post-pearling distancing obtained after 20-30 seconds of simulation, and pre-pearling distributions recapitulated after ~2 minutes (smallest Kolmogorov-Smirnov D statistic to experimental distributions). It is important to note that these are lower bounds for the values, as published nucleoid velocities do not account for mitochondrial movement ^6^. Nonetheless, given that pearling establishes regular distancing of nucleoids, intra-mitochondrial diffusion and nucleoid aggregation are sufficient to explain experimental post- and pre-pearling distributions.

Taken together, our data provide evidence that pearling establishes an emergent length scale, characteristic to the organelle, which dynamically entrains nucleoids sufficiently frequently to maintain their distribution across the network.

### Pearling onset drives nucleoid disaggregation

Nucleoid intensity profiles became smaller and sharper following pearling onset, with the full-width half-maxima of nucleoids (Fig. 3a) consistently decreased after pearling, trending towards their expected diffraction-limited size (377 ± 61 nm post-pearling). While nucleoid compaction could lead to smaller sizes, their typical ~100-200 nm diameter^32,33^ is beyond the resolution achievable in live-cell compatible microscopy methods, resulting in nucleoids appearing larger than they are (Fig. 3b). STED imaging reveals that each nucleoid consists of an average of ~1.6 DNA puncta each, both in U2-OS and HeLa cells (1.64 ± 0.12 and 1.60 ± 0.18 respectively), consistent with estimates of mtDNA copy number per nucleoid^33^. While ~57% of nucleoids contain a single DNA punctum, another ~40% contain 2 or 3 STED-resolved puncta (Fig. 3c). The dispersion of multi-copy nucleoids via pearling into single-copy smaller nucleoids may explain the decrease in size and average inter-nucleoid distances upon pearling (Fig. 3d).

**Figure 3.**
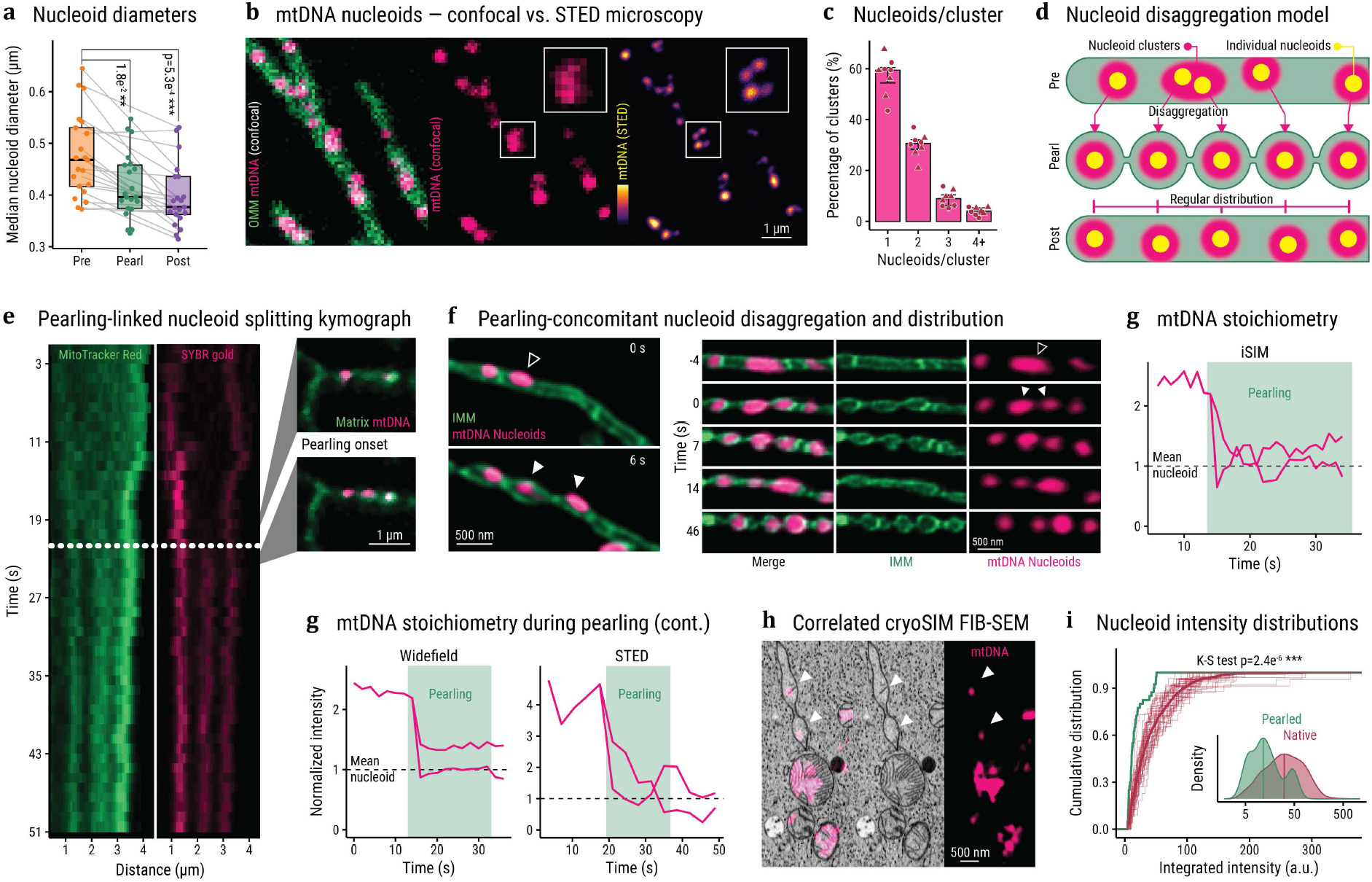
Pearling mediates nucleoid disaggregation. **(a)** Median nucleoid diameter (FWHM of TFAM-RFP peak) per independent event during pearling (Pearl), as well as within 10s before and after (Pre and Post). Boxplots represent median and IQR, p values from a two-sided Mann-Whitney test (n = 21 independent events with an average of 5.5 nucleoids and 15 time points each). **(b)** Immunofluorescent images of mtDNA nucleoids labeled by an anti-dsDNA antibody using confocal (pink) and STED (inferno) imaging, as well as confocal imaging of mitochondria (anti-TOMM20, green). **(c)** Percentage of nucleoid clusters segmented from confocal imaging as in (b), according to the number of STED-resolved nucleoids in U2-OS and HeLa cells (circles and triangles respectively). **(d)** Working model of pearling-mediated nucleoid splitting and redistribution (mitochondria in green, nucleoid clusters in pink, individual nucleoids in yellow. **(e)** Kymograph of nucleoid disaggregation concomitant with pearling onset (dashed line), with snapshot insets 2 s before and after, iSIM imaging. **(f)** Nucleoid segregation events from live STED imaging of IMM (PKMO, green) and mtDNA (SYBR gold, pink) during spontaneous pearling. Arrowheads indicate nucleoid splitting (filled, pre; empty, post). **(g)** Integrated intensity of nucleoids splitting during spontaneous pearling from different microscopes, normalized to the mean of neighboring non-splitting nucleoids (dashed line). Pearling duration shaded in green. **(h)** Spontaneous pearling FIB-SEM correlated with cryoSIM of nucleoids (SYBR green stain, pink). **(i)** Cumulative distribution of nucleoid integrated intensity from data as (h), manually classified by their presence within pearled mitochondria in FIB-SEM (n = 1061+40 in native and pearled mitochondria respectively). Thick lines represent the whole population (p-value from a Kolmogorov-Smirnov test), thin lines represent the bootstrapping of 40-nucleoid groups, inset with log_10_ intensity distributions. IMM = inner mitochondrial membrane, mtDNA = mitochondrial DNA.

Indeed, we captured splitting events concomitant with pearling (Fig. 3e-f, Supplementary Movie 2), resulting in the separation of a larger and brighter nucleoid focus into two smaller, dimmer, and more spherical foci. The stoichiometry of SYBR gold fluorescence intensity before and after splitting indicates the nearly equal separation of mtDNA material into the two resulting nucleoids (Fig. 3g). As an independent measure, we segmented and quantified nucleoids from cryo-preserved correlative whole-cell FIB-SEM and volumetric SIM imaging (Fig. 3h), classifying them by the pearling status of their mitochondrion in the electron micrograph. Nucleoids within pearled mitochondria presented significantly lower integrated SYBR green intensity than the native population average (Fig. 3i, Extended Data Fig. 2e-f), indicating they contain fewer mtDNA copies.

Pearling frequency data indicated that up to 11% of the total network length may pearl within a minute in control conditions, typically multiple times. Assuming all mitochondria are equally likely to pearl (Extended Data Fig. 1a), every part of the network can pearl multiple hundreds of times within a cell cycle. With the quantified 1.5 ± 0.4 pearling events/cell/min under gentle imaging conditions, we can estimate a lower bound of 2,610 pearling events per 29 h U2-OS cell cycle, or about 14,000 individual constrictions when factoring in the average number of pearls per event. This would be largely sufficient to segregate the duplication of each of the ~1000 nucleoids typically contained in a U2-OS cell. These rough approximations indicate that steady-state pearling frequencies are likely sufficient to disaggregate newly made mtDNA copies and maintain nucleoid separation along the network throughout cellular doubling time.

### Nucleoid distance regularity is independent of mitochondrial fission and membrane tethering

The inhibition of mitochondrial fission (typically by removal of its executor GTPase DRP1) was reported to increase pearling frequency and duration^21,23,25^. However, the modulation of proteins key to mitochondrial dynamics frequently leads to the loss of normal mitochondrial morphology (swelling, fragmentation), confounding interpretations of causality in their consequences on nucleoid morphology^16^. To circumvent off-target effects and forego severe pathology, we silenced the mitochondria-specific DRP1 adaptor proteins mediating each type of mitochondrial fission (proliferative midzone fissions by siMFF and damage-linked peripheral fissions by siFIS1), along with the inner membrane scaffolding and nucleoid tethering protein ATAD3A^34^. The inhibition of either type of fission indeed led to hyperfused mitochondrial networks (Extended Data Fig. 2g-i) and increased probability of pearling (Fig. 4a). In contrast, the downregulation of both ATAD3 isoforms did not affect pearling frequency (Fig. 4a), and maintained network structure and nucleoid dispersal (Extended Data Fig. 3k-l) as previously reported^17^, suggesting that nucleoid and IMM-OMM tethering are dispensable for pearling and pearling-driven nucleoid segregation.

**Figure 4.**
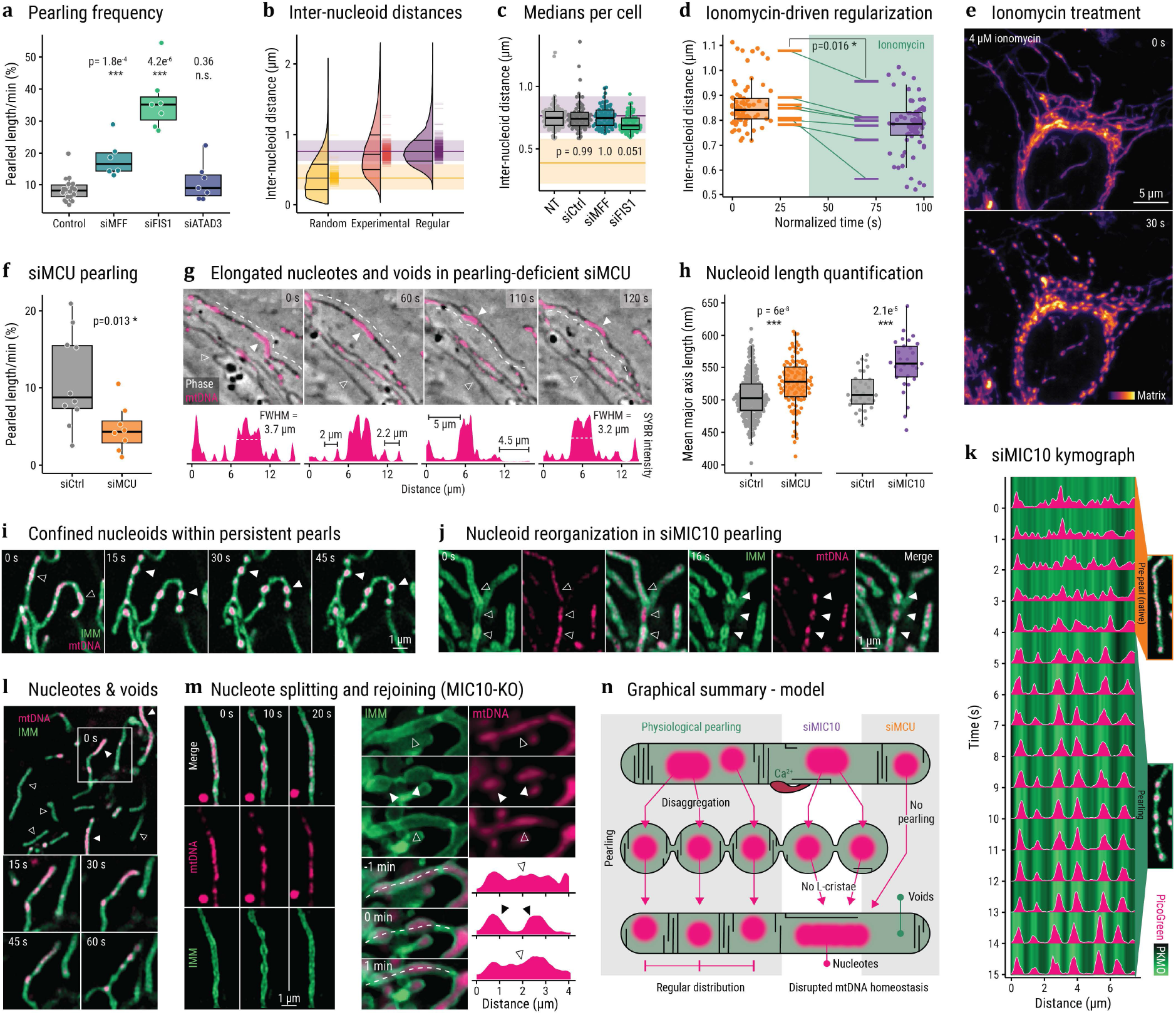
Modulating pearling affects nucleoid distribution. **(a)** Pearling frequency across siRNA treatments, percentage of the total mitochondrial network length pearling per minute (n = 18+6+7+7). p values from two-sided Wilcoxon tests. **(b)** Distribution of 3D inter-nucleoid distances pooled among siMFF and siFIS1 experiments (Experimental), along with models (Random and Regular). Horizontal bars represent median and IQR, and barcode plot depicts median values per cell (n =72953 nucleoids across 453 cells, 2 independent experiments). **(c)** Median inter-nucleoid distance per cell in control or fission-deficient cells. Horizontal shaded lines indicate median and IQR of random and regular distribution models as in (b), p values from an ANOVA/Tukey’s HSD test against non-treated cells (n = 87+69+71+60 cells across 2 independent experiments). **(d)** 3D inter-nucleoid distances before and during ionomycin-induced pearling. Points indicate median inter-nucleoid distance over time (centered around pearling onset), summarized by the boxplots, barcode plots represent the median value per independent experiment, p-value from a paired two-sided Student’s t-test (n = 12 independent experiments). **(e)** Maximum intensity projection of a U2-OS cell mitochondrial network undergoing ionomycin-induced pearling (light sheet imaging) (n = 12 independent experiments). **(f)** Pearling frequency in control and MCU-silenced cells (n = 10+8 cells), quantified as in (a). **(g)** Timelapse of mitochondrial and nucleoid dynamics (dark tubules in phase-contrast and SYBR gold in pink, respectively) in MCU-deficient cells (n = 15 control + 18 siMCU cells, across 2 independent experiments). Filled and empty arrowheads indicate a nucleote a nucleoid-free mitochondrial stretch respectively. Below, intensity line profiles of SYBR gold fluorescence along the dashed line (displaced for clarity) with key measures (nucleote sizes, inter-nucleoid distances, nucleoid void stretches). **(h)** Quantification of average nucleoid length per cell. Each point indicates the mean major axis length of all segmented nucleoids in a cell, p values from two-sided Wilcoxon tests. **(i)** Persistent spontaneous pearling event in siMIC10 HeLa cell, labelling IMM (PKMO, green) and mtDNA (PicoGreen, pink). **(j)** Reversible pearling event as in (i), highlighting nucleoid organization before (empty arrowheads) and during (filled arrowheads) pearling. **(k)** Kymograph of a spontaneous siMIC10 pearling event, mitochondrial (PKMO) signal overlaid with line profile of mtDNA (PicoGreen), pre- and pearling periods indicated with snapshot insets. **(l)** Overview of siMIC10 nucleoid defects: nucleotes (filled arrowheads) and nucleoid-free mitochondria stretches (empty arrowheads), and timelapse of pearling-mediated nucleote disaggregation as inset. **(m)** Confocal (left) and STED (right) imaging of cristae (PKMO, green) and nucleoids (PicoGreen, pink) in MIC10-KO HeLa cells, exemplifying separation and rejoining of nucleotes around pearling events. Line profile of PicoGreen intensity along the dashed line as inset. **(n)** Graphical model of pearling-mediated nucleoid reorganization in physiology and modulation conditions: Physiological mitochondria (green) undergo pearling (likely triggered by ER-derived calcium, in red) and recover, thus disaggregating enlarged nucleoids, establishing regular distancing, and maintaining separation by lamellar (L-) cristae. The inability to trigger pearling (siMCU) or maintain nucleoid separation because of lamellar cristae loss (siMIC10) leads to leads to nucleoid aggregation into nucleotes and nucleoid voids. Box plots represent median ± IQR of the overlaid datapoints. IMM = inner mitochondrial membrane, dsDNA = double-stranded DNA, NT = non-treated, si- = siRNA-silenced, siCtrl = siRNA non-targeting control.

Using segmented and skeletonized whole mitochondrial networks from volumetric iSIM immunofluorescent images, we quantified the distance along the 3D network between mtDNA intensity peaks. Then, using the total length and nucleoid number of each mitochondrion, we modeled two limit cases of nucleoid distributions: perfectly equidistant, or randomly distributed. As expected, the experimental distribution of distances was non-random, peaking at a median value very near to that of precise equidistance per mitochondrion (789 ± 425 and 808 ± 312 nm respectively), albeit broader (Fig. 4b-c). Our analysis points to a limiting factor in achieving equidistant nucleoid spacing across all mitochondria in a cell as the quantized nature of pearls and nucleoids themselves: mitochondrial lengths take on a continuous distribution, which prevents them from all partitioning into identical discrete units.

The inhibition of fission did not affect the inter-nucleoid distance; instead, both the median value and variability coefficient were maintained (Fig. 4c, Extended Data Fig. 2j). This indicates that normal nucleoid distribution is independent of fission dynamics, consistent with reports in yeast^18^. Interestingly, the proximity of steady-state distancing to precise equidistance and its insensitivity to increased pearling occurrence suggests that physiological inter-nucleoid distributions are maintained near the limit of precision achievable via this mechanism.

### Pearling-driven nucleoid distribution is triggered by intra-mitochondrial calcium and maintained by cristae confinement

We sought to identify regulators of pearling-driven nucleoid redistribution through additional manipulations. Ionomycin, a calcium ionophore, can induce simultaneous pearling across the entire mitochondrial network^24,25^, typically within 15-30 seconds of addition (Fig. 4d-e). We used ionomycin to controllably trigger widespread induction of pearling and dynamically measure its effect on nucleoids. We consistently found a reduction in cell-wide mean distances and regularization of the three-dimensional inter-nucleoid distance distribution compared to pre-treatment (Fig. 4d), supporting a causal relationship between network pearling and nucleoid distribution.

Since calcium signaling can trigger pearling^25^, we performed siRNA-mediated depletion of the mitochondrial calcium uniporter (siMCU). As expected, this decreased the probability of pearling (Fig. 4f, Extended Data Fig. 2m-n) with little consequence on mitochondrial morphology. While sections of the network presented normal nucleoid populations, we consistently observed enlarged and elongated nucleoids (over 2 μm long), hereby named ‘nucleotes’ (/*nu*.*kle ′o*.*tes*/), along with long stretches of nucleoid-free mitochondria (Fig. 4g). The presence of nucleotes was also observable in the significantly increased per-cell mean nucleoid major axis length in siMCU cells (Fig. 4h). Nucleotes were typically stable during ~1.5 min acquisitions; apparent movement or bending from mitochondrial branching rarely resulted in their disaggregation. The disruption of nucleoid morphology and coverage in pearling-deficient siMCU cells supports the importance of intra-mitochondrial calcium in triggering pearling to drive nucleoid segregation.

In wild-type cells, nucleoids typically reside in regions devoid of cristae (Extended Data Fig. 2o, ^28^). The exceptional pearls lacking nucleoids frequently presented high IMM signal, suggesting cristae density may sterically exclude nucleoids (Extended Data Fig. 2a). Thus, we sought to dissect the role of cristae in nucleoid distribution, taking advantage of siMIC10 as a model for lamellar cristae loss and increased pearling probability and stability (Fig. 1f-h). Most persistent pearls contained a nucleoid focus (Fig. 4i), and we observed nucleoid splitting and redistribution concomitant with reversible pearling onset (Fig. 4j-k), similar to steady-state conditions. Despite the heightened pearling prevalence, tubular siMIC10 mitochondria frequently contained nucleotes up to 6 μm long and stretches of nucleoid-free mitochondria, morphologically similar to those in siMCU cells (Fig. 4h,l). Nucleote-containing mitochondria were particularly devoid of lamellar cristae (Extended Data Fig. 2p), and these phenotypes were replicated in a stable MIC10-KO cell line (Fig. 4m, Extended Data Fig. 2p)^30^.

Pearling in siMIC10 cells readily separated nucleotes into smaller foci (Fig. 4l-m); however, nucleotes divided by pearling in MIC10 deficient cells tend to rapidly remerge. This contrasts with wild-type cells where post-pearling mitochondrial tubules preserve nucleoid separation. This indicates that lamellar cristae integrity is required to retain inter-nucleoid distancing post-pearling. We conclude that either the inability to pearl, as in the siMCU condition, or to constrain nucleoid motion, as in the siMIC10 condition, can cause nucleoid clustering and defective mtDNA distribution (Fig. 4n).

Taken together, our data supports a model wherein nucleoid disaggregation and distancing is independent of mitochondrial fission or membrane tethering, and is mechanistically driven by calcium-triggered pearling. Furthermore, it emphasizes the critical importance of lamellar cristae integrity for the resistance to and recovery from pearling, as well as to maintain the established nucleoid separation.

## Discussion

Despite over a billion years of reductive co-evolution, mtDNA remains an indispensable organellar genome for most eukaryotic cells^5^. Whether it is due to the limited dispersal of the hydrophobic proteins they encode, the need for their accurate inheritance, or their ability to enable local rapid stress detection^2–6,15^, the presence of well-distributed mtDNA copies is fundamental for mitochondrial function. Defects in mitochondrial dynamics and mtDNA maintenance can cause mitochondrial diseases, severe human disorders with puzzling pathomechanism and no cure^1,10^. Disease phenotypes often exhibit disrupted mtDNA integrity, including point- and large-scale mutations and topological reorganizations, depletion, and defective nucleoid distribution^16,19,35–38^. Decreased mtDNA levels have additionally been linked to aging, neurodegeneration, and cancer^39^, but boosting copy number can result in nucleoid enlargement and pathological mitochondrial stress signaling^37,40^, emphasizing the importance of precise nucleoid coverage and homeostasis for cellular and organismal health.

Despite its fundamental significance and its first observation decades ago, the mechanism underlying the regular distribution and segregation of nucleoids has remained an open question^6,7^. The answer, we demonstrate, lies in the intermittent undulations of mitochondria known as pearling. Physiological pearling events across the network generate steric constraints which position nucleoids at regular intervals, establishing a characteristic length scale without requiring active transport. Pearling propagates across interconnected mitochondria, translating a local ERMCS-linked calcium influx trigger^24–26^ into whole-organelle nucleoid organization. A fission event often follows pearling^24,25^, likely facilitated by the assembly of fission machinery in one of the constrictions^41^. Pearling thus provides a central mechanism for the maintenance of nucleoid distribution, coherent with the diversity of processes surrounding ER-mitochondrial dynamics and mtDNA proliferation^14,24,25,42,43^.

We systematically dissected pearling-driven nucleoid segregation while preserving mitochondrial tubular morphology, and identified the synergistic roles of calcium influx and lamellar cristae in modulating pearling and ensuring mtDNA nucleoid organization. Abrogating either process resulted in nucleoid aggregation into elongated *nucleotes*, a reversible form of nucleoid clustering within tubular mitochondria, distinct from enlarged nucleoids or “mito-bulb” clusters observed in engorged mitochondria in different pathologies^19,37,40,44^. Our data revealed that MCU-mediated calcium influx into mitochondria (likely through ERMCS) is a key inducer for pearling, consistent with published data^25^ and providing a mechanism to its recently reported association with nucleoid morphology in a mitochondrial-targeted siRNA screen^45^. In turn, lamellar cristae integrity (maintained by MIC10) plays a dual role: maintaining mitochondrial tubular stability and recovery from pearling, likely through enhancing its elasticity^20,29^, and confining newly segregated nucleoids after pearling recovery to avoid re-merging. The close relationship between cristae density, metabolic status, and respiratory efficiency^46^ raises the intriguing possibility that pearling frequency could connect local respiratory function with mtDNA quality control in a positive feedback loop.

Mitochondrial evolution into an elongated and dynamic network allows for metabolic and signaling plasticity, genetic complementation, and access to the cell’s farthest reaches^1,11^. However, in the absence of a cell wall, such tubular shapes under tension are subject to physical laws that can drive shape changes^8,47,48^. The pearling instability arises from the competition between elastic resistance and interfacial tension^20,26^, does not require energy consumption, and is reported in a wide variety of biological systems – long neuronal processes^47^, infected endosomal networks^49^, the ER^50^, and even bacterial substrate sensing and vesicle generation^51–54^. Our findings indicate that the localized occurrence of pearling is both common and readily reversible in physiological conditions, and is harnessed by the mitochondrial network to ensure the regular distribution of its genome.

## Methods

### Cell culture & live stains

Most experiments were performed in U-2 OS cells (ECACC 92022711) obtained from ECACC, which were grown in McCoy’s 5A Modified Medium (Gibco) supplemented with 10% fetal bovine serum, 100 U/ml penicillin and 100 mg/ml streptomycin. Other cell types used, including Cos-7 (ECACC 87021302) and HeLa cells (Cox8a-SNAP and MIC10 KO stable lines gift from T. Stephan and S. Jakobs ^30^) were grown in Dulbecco’s modified Eagle’s medium (DMEM) with similar supplementation. All cells were cultured in 5% CO_2_ at 37 °C and maintained in culture for a maximum of 20 passages. They were also routinely tested for *Mycoplasma* contamination and discarded if positive.

Plasmid transfection was achieved using Lipofectamine 2000 (Invitrogen) according to the manufacturer’s instructions, typically 2.5 μl of lipofectamine and 500 ng of DNA per 2 ml. Cells were imaged 12-48 h after transfection. All fluorescent protein constructs have been previously described. mt-StayGold (pCSII-EF/mt-(n1)StayGold) and ER-StayGold (pcDNA3/er-(n2)oxStayGold(c4)v2.0) were a gift from A. Miyawaki (Addgene plasmids #185823 and #186296), TFAM-RFP was derived from pMIH-TFAM-GFP plasmid, a gift from B. Kile (Addgene plasmid #113704).

siRNA treatments were designed to disrupt essential pathways only partially and acutely, aiming to forego severe pathology, loss of tubular morphology, swelling, or fragmentation ^30,55,56^, using published ^17,25,43^ and newly designed sequences. The transfection was performed using the manufacturer’s instructions of Lipofectamine RNAiMAX (Invitrogen), using 0.01-2 nmol of siRNA per ml of media and imaging cells 24-48 h after transfection. For siATAD3 and siMCU, cells were treated with a combination of several sequences. MIC10-targeted siRNAs were synthesized by siPOOLs Biotech (NCBI Gene ID 440574). Other oligonucleotides were synthesized by Microsynth: siMFF (5′-CGC UGA CCU GGA ACA AGG A-dTdT-3′), FIS1 (5′-CGA GCU GGU GUC UGU GGA G-dTdT-3′), siATAD3A (5’-GCA ACC AAC CAG AGC AGU UTT-3’), siATAD3B (5’-CCA AGG ACA AAU GGA GCA ATT-3’), siMCU_1 (5’-UAA UUG ACA CUU UAG AUU AUC UCU UTT-3’), siMCU_2 (5’-GUA ACA UAC UUC AUC ACU UAU GGA ATT-3’), siMCU_3 (5’-AUU GAC AGA GUU GCU AUC UAU UCA CTT-3’).

For dye-based stainings, the following reagents were used, following manufacturers’ staining instructions: Mitotracker Green (ThermoFisher M7514), Mitotracker Red FM (ThermoFisher M22425), Quant-iT PicoGreen (Life Technologies P7581), SYBR™ Gold Nucleic Acid Stain (Thermo Fisher Scientific, S11494), PK Mito Orange PKMO/ abberior LIVE ORANGE (Spirochrome SC053 /abberior LVORANGE), CellLight MitoRFP (Thermo, C10601).

W303 background-derived yFT016 *Saccharomyces cerevisiae* cells expressing Su9-mKate (mitochondria) were a gift from C. Osman and F. Thoma ^18^. Cells were grown in SCD and imaged under agarose pads.

Ethidium bromide treatment was performed at a concentration of 50ng/ml in growth media additionally supplemented with uridine and pyruvate over 48h, after which cells were stained and imaged. MtDNA quantification was done as described in the Image Analysis section.

### Fixation and immunofluorescence

For immunofluorescence, cells were plated on 12 mm glass coverslips (Menzel) 24 h before transfections or treatments. Cells were fixed with 3.2% paraformaldehyde (PFA) and 0.1% glutaraldehyde (GA) in phosphate-buffered saline (PBS), followed by washing and storage in PBS. To quench GA autofluorescence, the samples were treated with 0.5% freshly prepared sodium borohydride (NaBH_4_) for 7 minutes. Permeabilization and blocking were carried out using 3% bovine serum albumin (BSA) and 0.1% Triton X-100 in PBS for 45-60 minutes. Primary antibodies were incubated overnight in 1% BSA at 4°C. The samples were then washed twice for 10 minutes each with 1% BSA in PBS. Secondary antibodies were applied at room temperature for 1 hour in 1% BSA. The samples were subsequently washed twice with PBS and rinsed once with ddH_2_O, then immediately mounted using ProLong Gold Antifade Mountant (P36930, Invitrogen), allowing them to cure for 24 hours at room temperature in the dark. The antibodies used were dsDNA antibody (ab27156, Abcam) at a 1:250 dilution and TOM20 antibody (ab186734, Abcam) at a 1:200 dilution in 1% BSA, Alexa Fluor 488 anti-mouse, Alexa Fluor 568 anti-rabbit and Alexa Fluor 647 anti-mouse (A-11011, A-32723 and A-21235, Invitrogen), diluted 1:400 in 1% BSA.

### Light microscopy

Widefield imaging was performed on a Zeiss Axio Observer7 inverted microscope, equipped with an sCMOS camera (Photometrics Prime), a CoolLED pE-800 LED illumination system, and a 63x 1.40 NA oil immersion objective (Zeiss Plan-Apochromat 63/1.3 Oil Ph3 M27). Volumetric yeast data was acquired on a Nikon Spinning Disk CSU W1, fast confocal, 30 z planes acquired per timepoint.

Instant structured illumination microscopy (iSIM) was performed on a custom-built microscope set-up ^57^, equipped with 488-nm and 561-nm excitation lasers, a 1.49 NA oil immersion objective (APONXOTIRF; Olympus), an sCMOS camera (PrimeBSI, 01-PRIME-BSI-R-M-16-C; Photometric), and rapid vertically scanning piezo actuated mirrors for illumination pattern homogenization. Deconvolution of the raw images was done using the Richardson Lucy algorithm implementation in the flowdec Python package ^58^ with 10 iterations. Volumetric imaging was performed using a z-step size of 200 nm.

Array-detector confocal imaging was performed using (1) a Nikon AX-R laser scanning confocal microscope equipped with the NSPARC detector, using a Nikon Plan Apochromat 60X 1.42 NA oil immersion objective and processed on the fly in the Nikon NIS-Elements software, and (2) a Zeiss LSM 980 Axio Observer inverted microscope equipped with a Colibri 5 illumination system, an Airyscan 2 detector and a 63x 1.40 NA oil immersion objective (Zeiss Plan-Apochromat 63/1.4 Oil). Videos have a 1s frame rate for up to 360s, 40 nm pixel size, 0.65 μm pixel dwell time, and 2X SR sampling. Deconvolution and reassignation of the raw data were done using the Airyscan 2D SR Standard processing on the ZEN Blue software, version 3.4.91.

Single objective light-sheet imaging (SNOUTY, ^59^) was performed using a fixed Nikon 100x 1.35 NA silicon objective (MRD73950) with 488nm and 561nm excitation lasers and ZET405/488/561/640 quad filter wheel. All imaging was performed in a stage-top environmentally controlled incubator at 37C in 8-well glass-bottom chambered cover slides (CellVis C8-1.5H-N) pre-coated with 1:100 fibronectin (Sigma, F1141). Cells were imaged at 1 volume/second (1 Hz). At 30-45 seconds the well was injected with 2 μl of 400 μM ionomycin (Thermo, I24222) for a final concentration of 4 μM in the well. Cells were then imaged for a total of 200 volumes. The 10 frames pre-pearling were selected before the injection of ionomycin, and the 10 frames after were selected once pearl morphology started to become visible.

Live STED imaging was performed on an Olympus IX83 equipped with an Abberior STEDYCON system and a 100X 1.45 NA oil immersion objective (Olympus UPLXAPO100XO). STED laser was a pulsed 775 nm laser with a 40 MHz repetition rate, a max power of 1.25 W, and a pulse duration of 1.3 ns. The Pulsed Diode 488 and 561 lasers had a repetition rate of 40 MHz, and max power of 5 mW. For both channels, the excitation pulse delay was −0.2 ns, and gating was set for 1ns to 6 ns. For cristae imaging, a 561 nm laser was used for excitation, and emissions were detected in the range of 575-625 nm. For nucleoid imaging the excitation was with a 488 nm laser, and emissions were detected in the range of 505-550 nm. The pixel size was fixed to 30 nm x 30 nm. Pixel dwell time was 10 μs, 10 line accumulations. Pinhole was set to 0.91 AU. Frame duration was set to the minimal time allowed per field of view, resulting in frame rates as low as 3 seconds. Cristae data was deconvolved with Huygens Professional using the conservative fully automated smart template (Scientific Volume Imaging). 55 live STED acquisitions were collected across 13 separate samples (each sample was imaged up to 1h), imaged during 4 imaging sessions on separate days.

Correlative STED/Confocal microscopy was performed on a Leica SP8 STED 3X with a 100X 1.47 NA oil immersion objective (Leica HC PL APO CORR), a supercontinuum laser for excitation (561 and 647 nm) and 775 nm STED depletion. Images were acquired in confocal and STED mode subsequently, with pixel sizes of 0.0971 and 0.0116 μm respectively, and a dwell time of 1.5 μs, using the recommended detection ranges from the manufacturer’s instructions.

### Pearling-driven acquisition

Smart imaging of pearling in phase contrast images was performed following a hybrid-EDA paradigm and custom software built on the pymmcore-plus environment, as described in ^27^, on the Zeiss AxioObserver7 described above at 37°C and 5% CO_2_. Briefly, three consecutive frames of a time series at 1 Hz were supplied in the channel dimension to a custom-trained U-net of depth 2 with 16 initial filters. The model was trained on a total of 132 time series of phase contrast data with 66 labeled pearled mitochondria. A threshold was applied to the output event score map and fluorescence imaging was activated only following detection of pearling in phase contrast.

### Correlative cryo-SIM/FIB-SEM

For FIB-SEM and cryo-correlative fluorescence imaging, cells were high-pressure frozen to avoid the generation of fixation artifacts and to immobilize the cells rapidly enough to capture the highly transient pearled states. Cells were seeded on sapphire coverslips (3mm diameter, 50 m thickness, Nanjing Co-Energy Optical Crystal Co. Ltd, COE) prepared as described in ^60^. In brief, coverslips were washed in a basic piranha solution (5:1:1 water : ammonium hydroxide : hydrogen peroxide), washed in distilled water, sputter coated with a thin layer of gold (sputter coater Desk II, Denton Vacuum), rinsed several times in distilled water, and pre-coated with 100 μg of Matrigel (Corning) for 3 hours at 37 degrees.

U2-OS cells were seeded and allowed to grow for 18 hours, at which time cells were stained with 50 nM TMRM (ThermoFisher, T668) and SYBR Green I 10000X concentrate in DMSO (ThermoFisher S7563) at a 1:500,000 ratio for 15 minutes. Cells were then gently washed three times with 10 ml of complete medium, taking care not to flip over the coverslips. The cells were then given 15 minutes to equilibrate in 2 ml of complete medium to ensure TMRM staining was calibrated to the underlying membrane potential (membrane potential was later used to ensure the selected mitochondria were healthy and functional).

Immediately before freezing, cells were manually inspected using an inverted widefield microscope to ensure reasonable morphology and viability. Cells were then transferred to a water-jacketed incubator (ThermoFisher, Midi 40) and kept at 37 °C in 5% CO2 and 100% humidity until ready for freezing. Coverslips were removed one at a time from the incubator, overlaid with a 25% (w/v) solution of 40,000 MW dextran (Sigma), loaded between two hexadecane-coated freezing platelets (Technotrade International), and placed in the HPF holder. Freezing was performed using a Wohlwend Compact 2 high-pressure freezer, according to the manufacturer’s protocol. Frozen samples were stored under liquid nitrogen until cryo-fluorescence imaging or freeze substitution.

Cryo-fluorescence imaging of cells was performed using a custom-built cryo-structured illumination microscope (SIM) as described in ^60^. Briefly, sapphire coverslips were removed from the encasing platelets under liquid nitrogen in a custom preparation chamber and loaded into a custom-designed coverslip holder, which was then transferred to the cryo-stage of the microscope using a commercial cryogenic vacuum transfer process. Following SIM image acquisition and processing, samples were returned to liquid nitrogen for storage until subsequent freeze substitution and resin embedding.

Freeze-substitution was performed with a modified version of a previously described protocol ^60,61^. Briefly, frozen samples were transferred to cryotubes containing freeze-substitution media (2% OsO4, 0.1% Uranyl acetate, and 3% water in acetone) and placed in an automated freeze-substitution machine (AFS2, Leica Microsystems). The samples were then washed three times in anhydrous acetone and embedded in Eponate 12 (Ted Pella, Inc.). The sapphire coverslip was then removed, and the block was re-embedded in Durcapan (Sigma) resin for FIB-SEM imaging.

FIB-SEM was performed as described in ^62^. Briefly, a customized FIB-SEM using a Zeiss Capella FIB column fitted at 90 degrees on a Zeiss Merlin SEM was used to sequentially image and mill 8 nm layers from the Durcapan-embedded block. Milling steps were performed using a 15 nA gallium ion beam at 30 kV to generate two sequential 4 nm steps. Data was acquired at 500 kHz pixel−1 using a 2 nA electron beam at 1.0 kV landing energy with 8 nm xy resolution to generate isotropic voxels. Data sets were registered post-acquisition using a SIFT-based algorithm.

Instance segmentation of mitochondria from 3D FIB-SEM volumes was generated by ariadne.ai. CryoSIM and FIB-SEM stacks were registered using the BigWarp Fiji plugin ^63^. In brief, multiple landmarks were selected between TMRM-labeled mitochondria in SIM images and stained mitochondria in the FIB-SEM stack. Based on these landmarks, an affine transformation was used to register and fuse the SIM and FIB-SEM stacks.

### Image analysis

To account for the many variables involved in pearling frequency calculations (*i*.*e*. framerate of detection, amount of mitochondria, size of events), several approaches were utilized to quantify pearling frequency. Two of such strategies were the number of cells presenting at least a pearled mitochondrion at frame one of an acquisition, and events per unit of time, manually quantifying every pearling event with an onset during a timelapse regardless of their location, size, or duration. These data were also used to calculate pearling counts, and regional enrichment of pearling, by classifying them based on radial sectioning of a cell in three regions based on the measured axis lengths of cells: perinuclear (within 20 μm of the cellular center of mass), midzone (between 20 and 30 μm) and peripheral (beyond 30 μm).

To account for mitochondrial amount in the field-of-view and pearling magnitudes (length of the total pearling event), we additionally quantified pearling frequency as a percentage of the total mitochondrial network to pearl within a minute. For this, we threshold-segmented, skeletonized, and measured the total mitochondrial network length at frame zero (using built-in CellProfiler 4.2.8 ^64^ functions), and then manually highlighted the mitochondrial lengths that pearled within the acquisition (typically between 1 and 2 minutes at 1 FPS). The ratio between the skeleton length of mitochondria that pearled and the total network length normalized to one minute represents the pearling frequency in percentage length/min.

For two-dimensional analysis of nucleoid positioning, manually annotated 504 nm-thick smoothed line profiles along the mitochondrion were used to obtain the integrated intensities of each channel. Nucleoid centroids were identified as peaks in the nucleoid channel profile (SYBR gold or TFAM-RFP), and their distances were measured as their difference in position along the smoothed line length. Thick line profiles were similarly used to obtain the intensity data for kymographs (nucleoid distribution, splitting) and full-width half-maximum measurements (pearl and nucleoid size quantifications). Classification of pearling status was performed blinded to the nucleoid channel, ensuring the periodical pearling morphology on both the mitochondrial channel image and line profile. Modeling of random distribution was done by assigning the nucleoids on each profile random positions along the length, reordering them, and recalculating the inter-nucleoid distances.

Whole-cell nucleoid quantification was performed using built-in CellProfiler 4.2.5 ^64^ functions. Briefly, the nucleoid signal was segmented using three-class global Otsu thresholding (through the IdentifyPrimaryObjects module), and sizes, shapes, and intensities were measured by the MeasureObjectSizeShape and MeasureObjectIntensity modules. For FIB-SEM correlative data, nucleoids were classified from a manual mask created from the electron microscopy data, identifying pearled regions blinded to the nucleoid signal. For correlative confocal/STED nucleoid analysis, segmented STED nucleoids were assigned to parent nucleoids segmented from the aligned confocal image, obtaining the number of STED objects per confocal nucleoid, as well as an array of properties of each (sizes, shapes, intensities of both confocal and STED nucleoids (mean) for each confocal nucleoid).

Nucleotes (long nucleoid clusters) were defined as clusters of nucleoid signal (from SYBR gold or PicoGreen) longer than 2 μm long and stably clustered throughout the timelapse (typically > 1 min). Nucleotes were differentiated from other enlarged nucleoid formations (mito-bulbs or enlarged nucleoids ^37,40,44^) by their maintaining mitochondrial tubular integrity. When measuring whole-cell nucleoid length averages, a CellProfiler-based strategy was utilized as above, using the per-cell mean major axis length measurement from the MeasureObjectSizeShape module to identify population shifts in nucleoid elongation.

Mitochondrial DNA amount quantification following ethidium bromide treatment was similarly performed, segmenting mitochondria using the IdentifyPrimaryObjects module in CellProfiler, and quantifying mean intensity of SYBR gold within the mitochondrion (quantitatively proportional to DNA amount), to confirm specific loss of mitochondrial DNA in a large number of cells and circumvent unspecific nuclear dye incorporation (consequential of mtDNA loss).

For volumetric analysis of inter-nucleoid distancing, 3D iSIM images of immunolabeled mitochondrial network and DNA were processed in a custom pipeline. Shortly, mitochondrial networks were segmented and skeletonized using Nellie ^65^, obtaining labeled skeletons. Splines were then generated from branches (above 1.12 μm), using the SplineBox package ^66^ to fit cubic B-splines to the skeleton coordinates, and extended by 300 nm to capture the entire mitochondrial length and potential tip nucleoids. Then, normal planes (1 μm wide) were computed along the splines, extracting the integrated intensity of each channel, and outputting a line profile similar to those from 2D data with distance information along the spline. The line profile, inter-nucleoid distancing, and random distribution analysis were performed as above. The regular distribution values were achieved by dividing the total length of a branch between the first and last nucleoid position and dividing it by the number of nucleoids on that branch minus one.

Volumetric live SNOUTY (baseline distancing, ionomycin treatment) and Spinning disk data (yeast) were analyzed using similar approaches. The 3D skeletons of both the mitochondrial network and nucleoids were extracted using Nellie ^65^. Nucleoid positions were obtained using a centroid algorithm with a minimum distance threshold, and subsequently matched/projected onto the nearest points on the mitochondrial network skeleton. The shortest path traversal-distance algorithm was then used to calculate the inter-nucleoid distances between neighboring nucleoids.

### Nucleoid diffusion simulation

The post-pearling nucleoid diffusion simulation consisted of a tubular container of 10 μm length and 400 nm wide representing a mitochondrion, with point objects located at the center of the tube and every 810 nm, representing ideally equidistant nucleoids immediately following pearling recovery. Using a diffusion coefficient of 894 nm^2^/s estimated from published nucleoid mean squared displacement values ^17,31^, each simulation was run for 600 seconds, with timesteps of 0.01 s. If nucleoids were <120nm from one another, the objects were merged in the average position, simulating imaging resolution and nucleoid aggregation. Every 10 s of simulation, the horizontal (x-axis) distances between neighboring nucleoids were measured. The reported data combines the results obtained from 100 simulations. The comparison to experimental distribution was performed through pair-wise minimization of the Kolmogorov-Smirnov D statistic.

### Western blotting

Immunoblotting was done on cells following identical and parallel siRNA treatments as imaging experiments. The cell pellets were lysed in RIPA buffer (Sigma) supplemented with freshly added protease inhibitors (Sigma Aldrich 11836170001) on ice for 30 minutes. Following lysis, the lysates were centrifuged at 16,000xg for 10 minutes at 4 °C to separate the insoluble material. Protein concentrations were quantified using the Pierce BCA protein assay kit (Life Technologies, 23227), and equal protein amounts (30-50 μg per lane) were resolved on self-cast 7.5% or 15% SDS-PAGE gels. For immunoblotting, proteins were transferred onto nitrocellulose membranes (BioRad) via electrophoresis and probed with the appropriate primary antibodies, which were diluted in 5% non-fat dry milk in Tris-buffered saline with Tween 20 (TBST). The blots were then incubated with anti-rabbit or anti-mouse horseradish peroxidase (HRP)-conjugated secondary antibodies (GE Healthcare) and visualized using ECL (GE Healthcare). The antibodies used were against: MIC10 (Abcam, ab84969, kind gift from Veronica Eisner) diluted 1:1000, FIS1 (LuBioScience GmbH, 10956-1-AP), diluted 1:2000, MFF (Life Technologies, PA5-67357), diluted 1:1000), MCU (Abcam, ab219827) tubulin (Abcam, ab18251), diluted 1:2000. Densitometry was performed using FIJI ImageJ software. Uncropped western blots can be found in Extended Data Fig. 3.

### Experimental and statistical design

Statistical analyses and plotting were performed in R version 4.4.2 within the Visual Studio Code environment version 1.93.0. No statistical method was used to predetermine the sample size. Only technically failed data (out of focus, no detectable signal) or visually identifiable deterioration of cellular health was excluded from the analysis, and partial blinding and automation were applied whenever possible to minimize observer bias (e.g. choice and classification of pearling regions using mitochondrial markers blinded to the nucleoid channel, and automatic quantification of inter-nucleoid distances blinded to genotype).

When reasonable, all data points are clearly displayed in data visualization and represent separate biological measurements (typically individual cells), overlayed on boxplots representing the median, two quartiles (Q2 and Q3), and minima and maxima as a central line, box bounds, and whiskers respectively. Outliers are defined as being beyond 1.5 times the interquartile range. Statistical tests were performed on relevant biological units (e.g. median per cell instead of each inter-nucleoid distance separately) to avoid non-informative inflated significance.

## Data & code availability

Data and code supporting this study are publicly available on the online repository Zenodo (DOI: pending) or GitHub (Link: pending), and plasmids and cell lines are available on request from the corresponding authors. Uncropped western blots are provided in Extended Data Fig. 3.

## Acknowledgements

The authors wish to thank Hélène Perreten, Kyle Douglass, Sheda Ben Nejma, Giorgio Tortarolo, Julius Winter, Christian Zimmerli, Veronica Eisner, Austin Lefebvre, Quentin Kohler, Carolin Klose, and Asa Kalish for their invaluable technical assistance and inspiring discussions. We thank the EPFL Center for Imaging (especially Florian Aymanns) for their support with image analysis. The EPFL Laboratory of Experimental Biophysics (LEB), the MBL Physiology Course and the HFSP “MitoManiacs” are also thanked for providing an enriching environment for discovery in fundamental science. We thank Stefan Jakobs and Till Stephan for kindly sharing their HeLa cell models, and Gleb Shtengel, Harald Hess, Eric Betzig, and Shan Xu for their contributions in developing the cryo-SIM FIB-SEM technique.

## Funding

Human Frontier of Science Program (HSFP, RGP0038/2021) (S.M., W.M.)

European Research Council (CoG 819823, Piko) (S.M.) The Swiss National Science Foundation (SNSF, 10268) (S.M.)

The Swiss State Secretariat for Education, Research and Innovation (SERI) CLS-HSG Research Partnership (SCR0835124) (S.M.)

Agencia Nacional de Investigación y Desarrollo de Chile (Ph.D. Fellowship 21211363) (J.M.)

Swiss Government Excellence Research Fellowship (FCS 2023.0400) (J.M.).

## Author contributions

Conceptualization: J.C.L., S.M.

Methodology: J.C.L., M.D.L., J.M., G.S., C.J.O., A.S.M., W.S., D.H.

Investigation: J.C.L., M.D.L., J.M., G.S., R.J., C.J.O., A.S.M.

Visualization: J.C.L.

Writing – original draft: J.C.L., S.M.

Writing – review and editing: All authors

Project administration: S.M., J.C.L.

Funding acquisition: S.M.

## Supplementary Materials

**Movie S1**.

iSIM time-lapse of a U2-OS cell mitochondrion undergoing consecutive reversible pearling, and eventually fission. Mitochondrial matrix labeled with mt-StayGold. Arrowheads and inset highlight pearling events, and empty arrowheads indicate fission events.

**Movie S2**.

Live STED timelapse of mitochondrial inner membrane (PKMO, green) and nucleoid (SYBR gold, pink) during spontaneous pearling in a U2-OS cell. Pink arrowheads indicate nucleoid foci and green arrowheads lamellar cristae.

**Movie S3**.

Timelapses of mitochondrial inner membrane (PKMO, green) and nucleoid (PicoGreen, pink) during spontaneous pearling in MIC10-deficient HeLa cells. Empty arrowheads indicate nucleoid-free mitochondria, filled arrowheads pearling events (reversible in pink, persistent in green) and rounded pink arrowheads mark nucleote aggregates.

**Extended Data Fig. 1.**
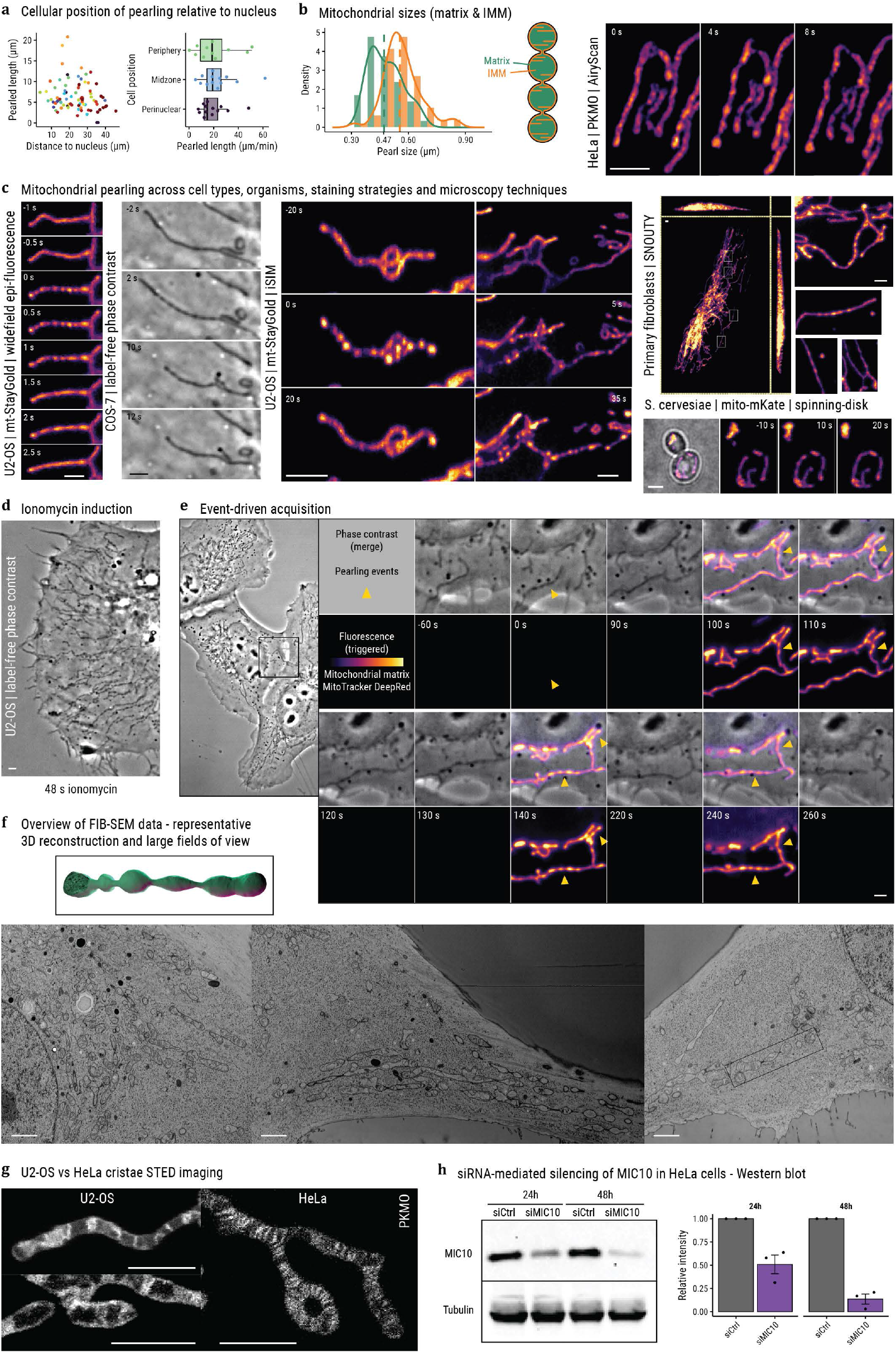
**(a)** Position of pearling events in the cell. Scatterplot of pearling event distance to the nucleus vs. its length (left, color indicates cell of origin), and pearling frequency classified by cellular region (right). **(b)** Distribution of pearl sizes (FWHM) as measured from matrix-targeted mt-StayGold on iSIM or IMM-targeted PKMO on STED. Dashed line indicates median value, a graphical depiction of different mitochondrial compartments on the right. **(c)** Montage of spontaneous pearling in a variety of models, contrasting methods, and microscopy techniques. Cell types: African green monkey kidney fibroblast-like COS-7, human osteosarcoma U2-OS, human cervical adenocarcinoma HeLa, Saccharomyces cerevisiae (budding yeast), and primary fibroblasts. Fluorescent markers: mt-StayGold, PK Mito Orange (PKMO), CellLight MitoRFP, and mt-mKate. Imaging was performed using widefield epi-fluorescent microscopy, label-free phase-contrast, instant structured illumination (iSIM), single-objective selective plane illumination (SNOUTY), and spinning disk confocal microscopy. **(d)** U2-OS cell following 48-second ionomycin treatment, label-free phase-contrast microscopy. **(e)** Pearling events detected during smart event-driven acquisition. Overview of the field-of-view (left) and highlighted timelapse of phase-contrast (top, constantly 1 FPS) and MitoTracker DeepRed fluorescence (bottom, only when triggered). Yellow arrowheads indicate pearling events, including one preceding any fluorescent imaging, and multiple detected by fluorescence. **(f)** Large fields of view of FIB-SEM in U2-OS cells, 3D reconstruction of a pearled mitochondrion on the right. **(g)** STED imaging of mitochondrial cristae (stained by PKMO) of wildtype U2-OS and HeLa cells. **(h)** Immunoblot of MIC10 siRNA efficiency and quantification, three repeats. All error bars represent 2 μm.

**Extended Data Fig. 2.**
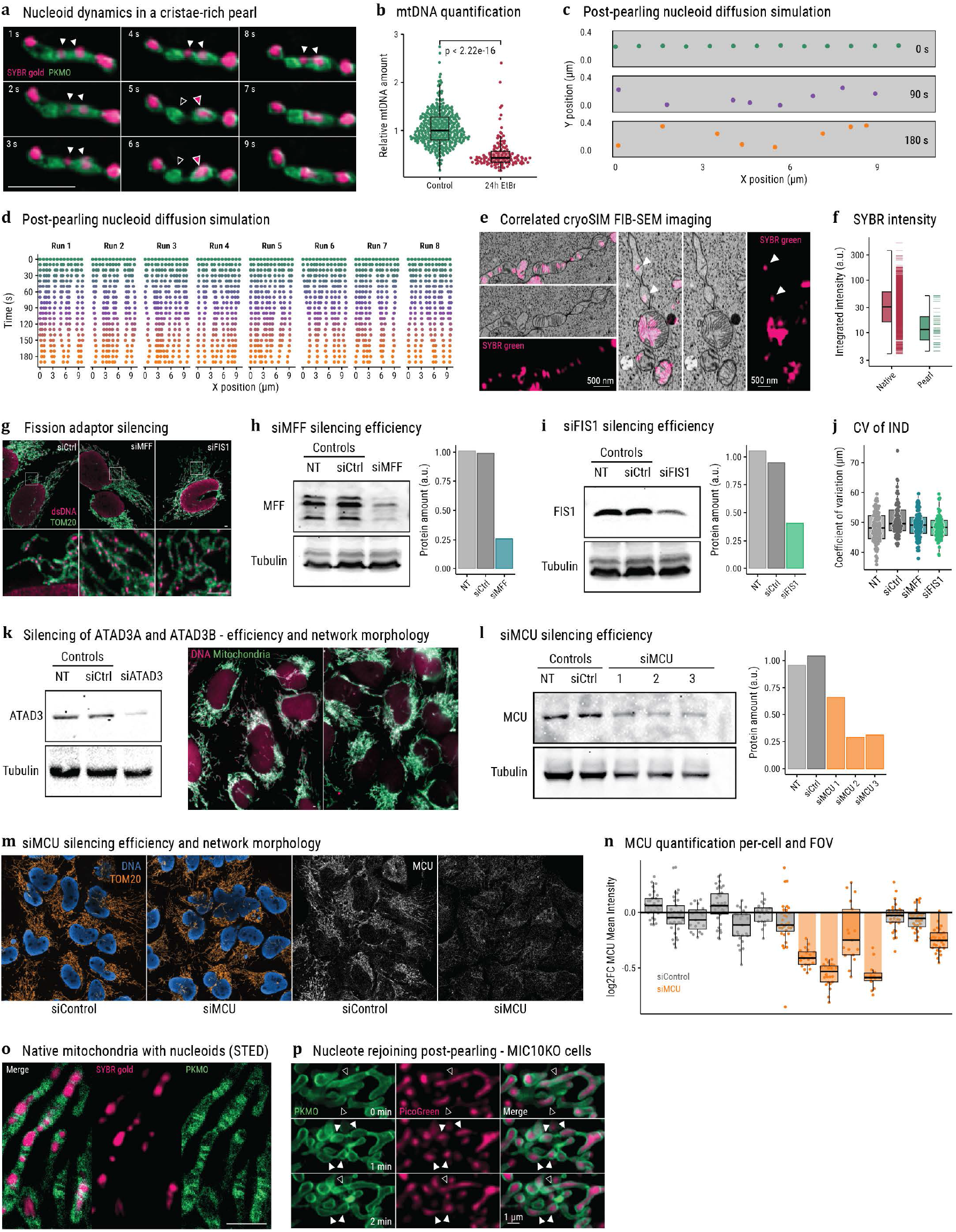
**(a)** Live STED timelapse of spontaneous pearling, mitochondrial IMM, and nucleoids stained with PKMO and SYBR gold respectively. Arrowheads highlight key pearls/nucleoids (white), as an IMM-rich pearl (empty arrowhead) loses its nucleoid to its neighboring pearl (pink arrowhead). **(b)** Quantification of mtDNA amount in ethidium bromide experiment, as mean SYBR gold intensity within segmented mitochondria relative to controls, p-value from two-sided Wilcoxon test. **(c-d)** Representative graphics of post-pearling nucleoid diffusion simulations: graphical depiction of the simulation tubular container (400 nm x 10 μm) with nucleoids at 0, 90, and 180 s time points (c) and 10 runs of the simulation only displaying nucleoids and their X position every 10 seconds. **(e)** Spontaneous pearling FIB-SEM correlated with cryoSIM of nucleoids (SYBR green stain, pink). **(f)** Distribution of nucleoid integrated intensity from data as (e), manually classified by their presence within pearled mitochondria in FIB-SEM (n = 1061+40 in native and pearled mitochondria respectively), y scale in log_10_ scale. **(g)** Volumetric immunofluorescence images of mitochondria and double-stranded DNA (TOM20 and dsDNA antibodies, max intensity projection), for control or silencing either MFF or FIS1 (n = 4 independent experiments). (**h-i)** Silencing efficiency of siRNA targeting MFF (h) and FIS1 (i), along with their densitometric quantification. **(j)** Coefficient of variation of 3D inter-nucleoid distances per cell in siRNA-treated cells. **(k-l)** Silencing efficiency of siRNA targeting ATAD3A+B (k) and MCU (l), along with their densitometric quantification. Three siMCU sequences were tested, which were combined for further experimentation. **(m-n)** Immunofluorescence of anti-TOM20, anti-dsDNA, and anti-MCU (orange, blue and white respectively) in siMCU (3 sequences combined) and non-targeting siRNA control cells (m), and quantification of mean MCU intensity per cell across replicates. All error bars represent 2 μm. **(o)** Output of the random forest model generated to predict STED-resolved nucleoid number as in Fig. 3c based on all size, shape, and intensity descriptors of segmented confocal objects, top 15 strongest predictors. **(p)** Live STED imaging of mitochondria and nucleoids (stained with PKMO and SYBR gold respectively. **(q)** Live STED imaging of cristae (PKMO) and nucleoids (PicoGreen) in MIC10-KO HeLa cells, highlighting nucleotes (empty arrowheads) separated into nucleoid-like foci upon pearling (filled arrowheads) and rejoined into nucleotes following recovery.

**Extended Data Fig. 3.**
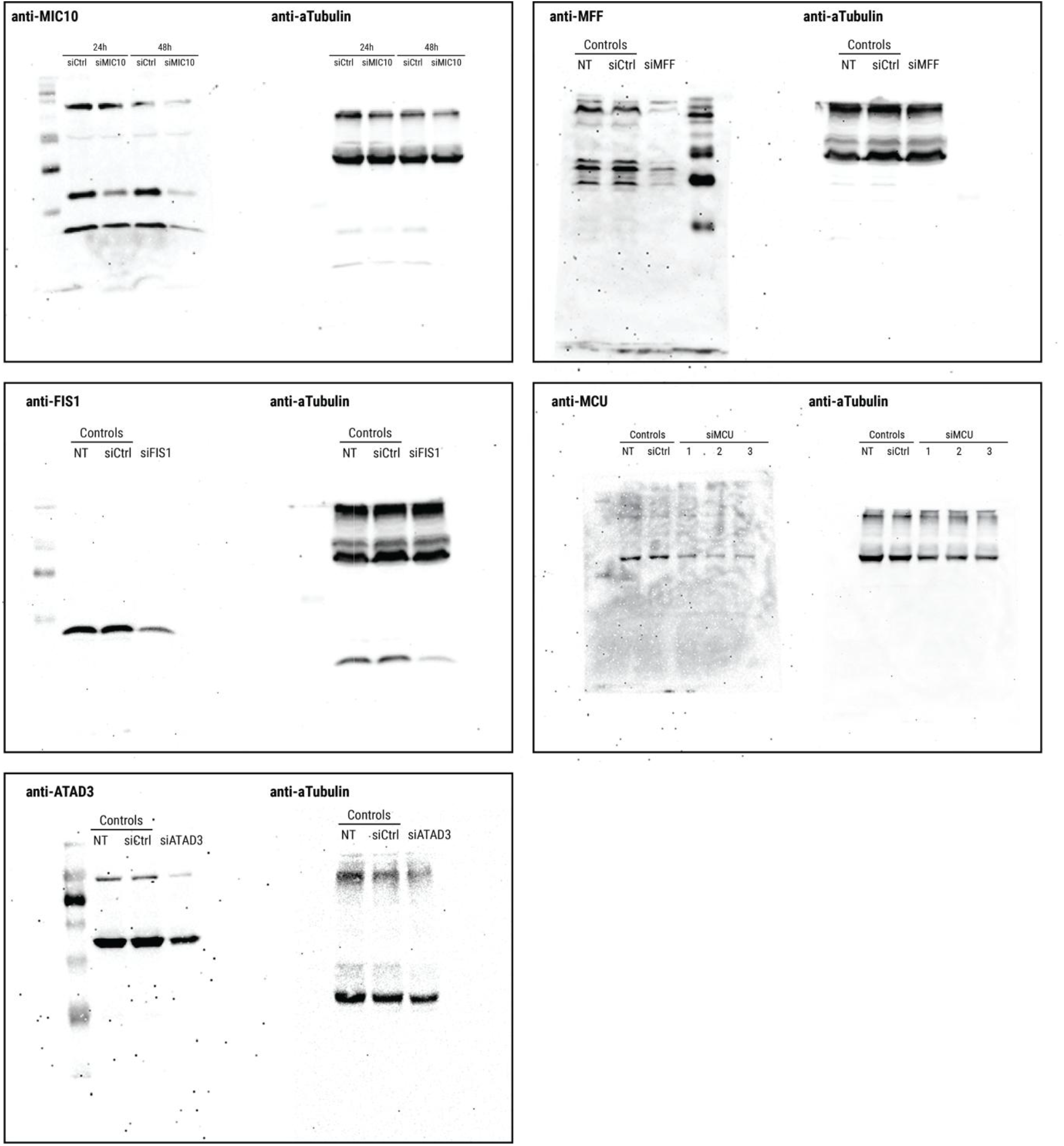
Uncropped western blots depicted in Extended Data Fig. 1-2, along with their loading controls.

