## Extended Data Figures for "Pearling Drives Mitochondrial DNA Nucleoid Distribution"

bioRxiv

### Movie S1.

iSIM time-lapse of a U2-OS cell mitochondrion undergoing consecutive reversible pearling, and eventually fission. Mitochondrial matrix labeled with mt-StayGold. Arrowheads and inset highlight pearling events, and empty arrowheads indicate fission events.

### Movie S2.

Live STED timelapse of mitochondrial inner membrane (PKMO, green) and nucleoid (SYBR gold, pink) during spontaneous pearling in a U2-OS cell. Pink arrowheads indicate nucleoid foci and green arrowheads lamellar cristae.

### Movie S3.

Timelapses of mitochondrial inner membrane (PKMO, green) and nucleoid (PicoGreen, pink) during spontaneous pearling in MIC10-deficient HeLa cells. Empty arrowheads indicate nucleoid-free mitochondria, filled arrowheads pearling events (reversible in pink, persistent in green) and rounded pink arrowheads mark nucleote aggregates.

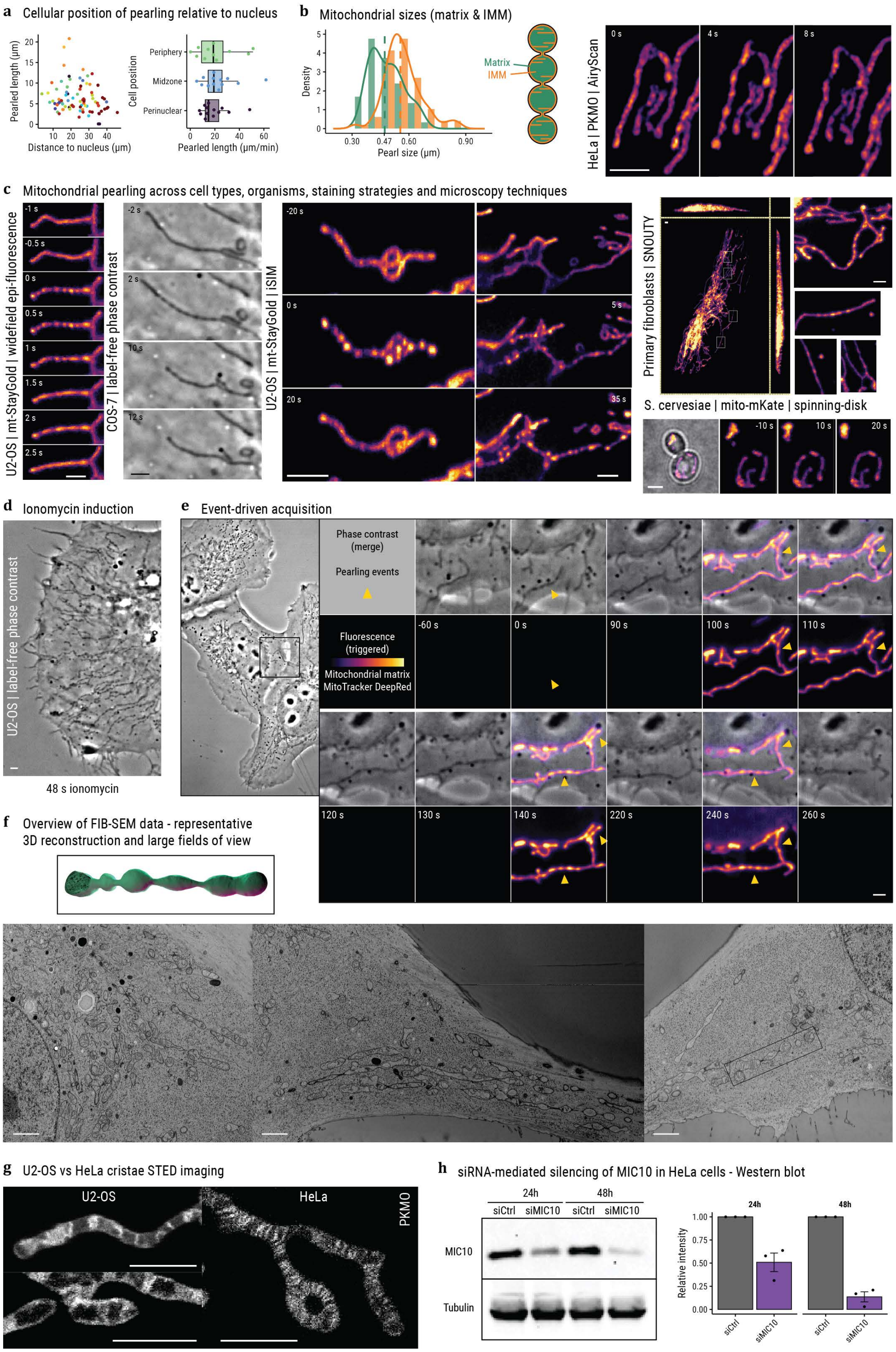
a
Cellular position of pearling relative to nucleus
Pearled length (µm)
Distance to nucleus (µm)
Cell position
Periphery
Midzone
Perinuclear
Pearled length (µm/min)
b
Mitochondrial sizes (matrix & IMM)
Density
Pearl size (µm)
Matrix
IMM
HeLa | PKMO | AiryScan
0 s
4 s
8 s
c
Mitochondrial pearling across cell types, organisms, staining strategies and microscopy techniques
U2-OS | mt-StayGold | widefield epi-fluorescence
COS-7 | label-free phase contrast
U2-OS | mt-StayGold | iSIM
Primary fibroblasts | SNOUTY
S. cervesiae | mito-mKate | spinning-disk
d
Ionomycin induction
U2-OS | label-free phase contrast
48 s ionomycin
e
Event-driven acquisition
Phase contrast
(merge)
Pearling events
Fluorescence
(triggered)
Mitochondrial matrix
MitoTracker DeepRed
-60 s
0 s
90 s
100 s
110 s
120 s
130 s
140 s
220 s
240 s
260 s
f
Overview of FIB-SEM data - representative
3D reconstruction and large fields of view
g
U2-OS vs HeLa cristae STED imaging
U2-OS
HeLa
PKMO
h
siRNA-mediated silencing of MIC10 in HeLa cells - Western blot
24h
48h
siCtrl
siMIC10
MIC10
Tubulin
Relative intensity
siCtrl
siMIC10

### Extended Data Fig. 1.

**(a)** Position of pearling events in the cell. Scatterplot of pearling event distance to the nucleus vs. its length (left, color indicates cell of origin), and pearling frequency classified by cellular region (right). **(b)** Distribution of pearl sizes (FWHM) as measured from matrix-targeted mt-StayGold on iSIM or IMM-targeted PKMO on STED. Dashed line indicates median value, a graphical depiction of different mitochondrial compartments on the right. **(c)** Montage of spontaneous pearling in a variety of models, contrasting methods, and microscopy techniques. Cell types: African green monkey kidney fibroblast-like COS-7, human osteosarcoma U2-OS, human cervical adenocarcinoma HeLa, Saccharomyces cerevisiae (budding yeast), and primary fibroblasts. Fluorescent markers: mt-StayGold, PK Mito Orange (PKMO), CellLight MitoRFP, and mt-mKate. Imaging was performed using widefield epi-fluorescent microscopy, label-free phase-contrast, instant structured illumination (iSIM), single-objective selective plane illumination (SNOUTY), and spinning disk confocal microscopy. **(d)** U2-OS cell following 48-second ionomycin treatment, label-free phase-contrast microscopy. **(e)** Pearling events detected during smart event-driven acquisition. Overview of the field-of-view (left) and highlighted timelapse of phase-contrast (top, constantly 1 FPS) and MitoTracker DeepRed fluorescence (bottom, only when triggered). Yellow arrowheads indicate pearling events, including one preceding any fluorescent imaging, and multiple detected by fluorescence. **(f)** Large fields of view of FIB-SEM in U2-OS cells, 3D reconstruction of a pearled mitochondrion on the right. **(g)** STED imaging of mitochondrial cristae (stained by PKMO) of wildtype U2-OS and HeLa cells. **(h)** Immunoblot of MIC10 siRNA efficiency and quantification, three repeats. All error bars represent 2 µm.

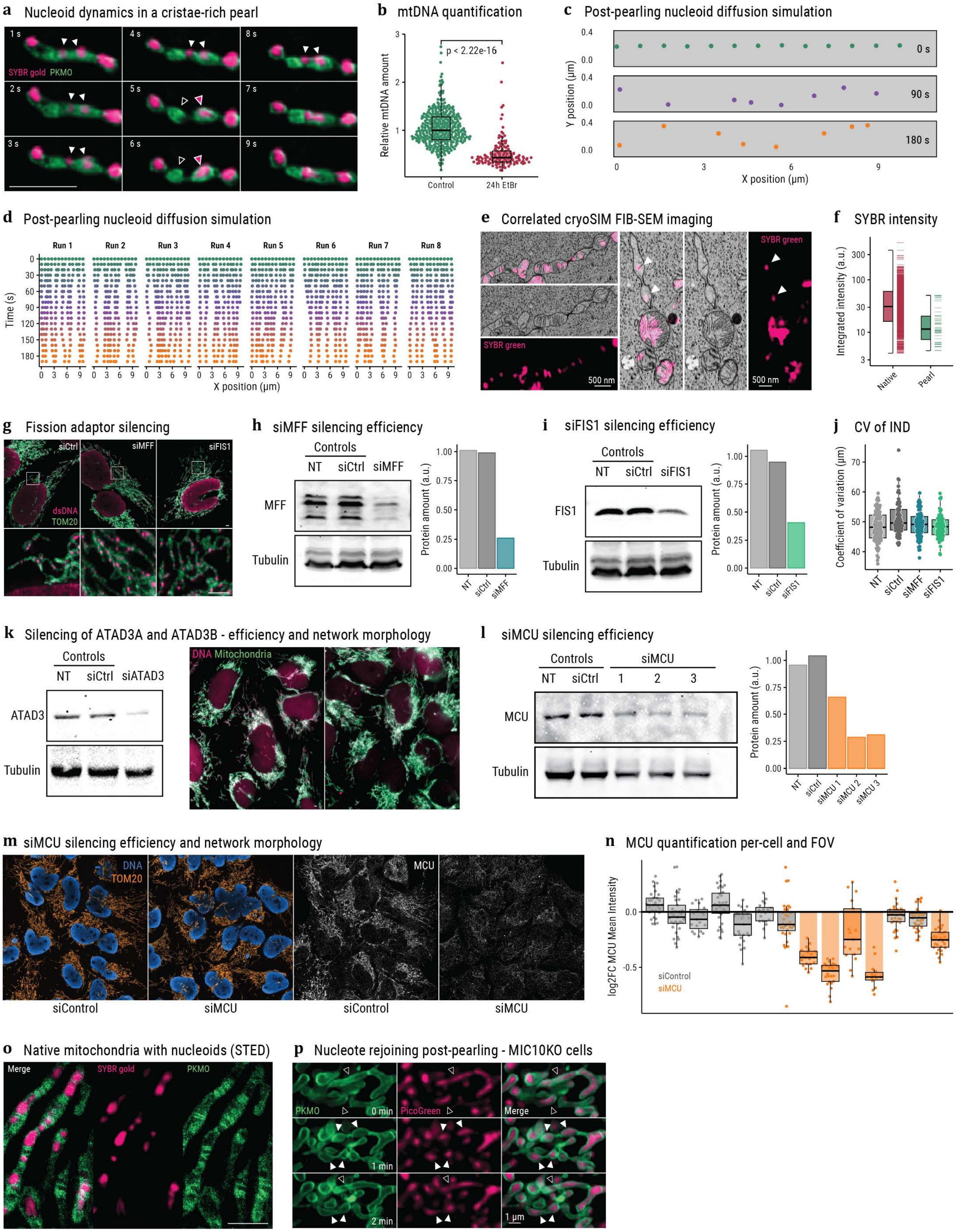

a Nucleoid dynamics in a cristae-rich pearl
SYBR gold PKMO
b mtDNA quantification
p < 2.22e-16
Relative mtDNA amount
Control
24h EtBr
c Post-pearling nucleoid diffusion simulation
0 s
90 s
180 s
Y position (µm)
X position (µm)
d Post-pearling nucleoid diffusion simulation
Run 1
Run 2
Run 3
Run 4
Run 5
Run 6
Run 7
Run 8
Time (s)
X position (µm)
e Correlated cryoSIM FIB-SEM imaging
SYBR green
500 nm
f SYBR intensity
Integrated intensity (a.u.)
Native
Pearl
g Fission adaptor silencing
siCtrl
siMFF
siFIS1
dsDNA
TOM20
h siMFF silencing efficiency
Controls
NT
siCtrl
siMFF
MFF
Tubulin
Protein amount (a.u.)
i siFIS1 silencing efficiency
siFIS1
FIS1
j CV of IND
Coefficient of variation (µm)
k Silencing of ATAD3A and ATAD3B - efficiency and network morphology
siATAD3
ATAD3
DNA Mitochondria
l siMCU silencing efficiency
siMCU
MCU
siMCU 1
siMCU 2
siMCU 3
m siMCU silencing efficiency and network morphology
DNA
TOM20
siControl
n MCU quantification per-cell and FOV
log2FC MCU Mean Intensity
o Native mitochondria with nucleoids (STED)
Merge
SYBR gold
PKMO
p Nucleote rejoining post-pearling - MIC10KO cells
PicoGreen
0 min
1 min
2 min
1 µm

**Extended Data Fig. 2.**

**(a)** Live STED timelapse of spontaneous pearling, mitochondrial IMM, and nucleoids stained with PKMO and SYBR gold respectively. Arrowheads highlight key pearls/nucleoids (white), as an IMM-rich pearl (empty arrowhead) loses its nucleoid to its neighboring pearl (pink arrowhead). **(b)** Quantification of mtDNA amount in ethidium bromide experiment, as mean SYBR gold intensity within segmented mitochondria relative to controls, p-value from two-sided Wilcoxon test. **(c-d)** Representative graphics of post-pearling nucleoid diffusion simulations: graphical depiction of the simulation tubular container (400 nm x 10 µm) with nucleoids at 0, 90, and 180 s time points (c) and 10 runs of the simulation only displaying nucleoids and their X position every 10 seconds. **(e)** Spontaneous pearling FIB-SEM correlated with cryoSIM of nucleoids (SYBR green stain, pink). **(f)** Distribution of nucleoid integrated intensity from data as (e), manually classified by their presence within pearled mitochondria in FIB-SEM (n = 1061+40 in native and pearled mitochondria respectively), y scale in $\log_{10}$ scale. **(g)** Volumetric immunofluorescence images of mitochondria and double-stranded DNA (TOM20 and dsDNA antibodies, max intensity projection), for control or silencing either MFF or FIS1 (n = 4 independent experiments). (**h-i)** Silencing efficiency of siRNA targeting MFF (h) and FIS1 (i), along with their densitometric quantification. **(j)** Coefficient of variation of 3D inter-nucleoid distances per cell in siRNA-treated cells. **(k-l)** Silencing efficiency of siRNA targeting ATAD3A+B (k) and MCU (l), along with their densitometric quantification. Three siMCU sequences were tested, which were combined for further experimentation. **(m-n)** Immunofluorescence of anti-TOM20, anti-dsDNA, and anti-MCU (orange, blue and white respectively) in siMCU (3 sequences combined) and non-targeting siRNA control cells (m), and quantification of mean MCU intensity per cell across replicates. All error bars represent 2 µm. **(o)** Output of the random forest model generated to predict STED-resolved nucleoid number as in Fig. 3c based on all size, shape, and intensity descriptors of segmented confocal objects, top 15 strongest predictors. **(p)** Live STED imaging of mitochondria and nucleoids (stained with PKMO and SYBR gold respectively. **(q)** Live STED imaging of cristae (PKMO) and nucleoids (PicoGreen) in MIC10-KO HeLa cells, highlighting nucleotes (empty arrowheads) separated into nucleoid-like foci upon pearling (filled arrowheads) and rejoined into nucleotes following recovery.

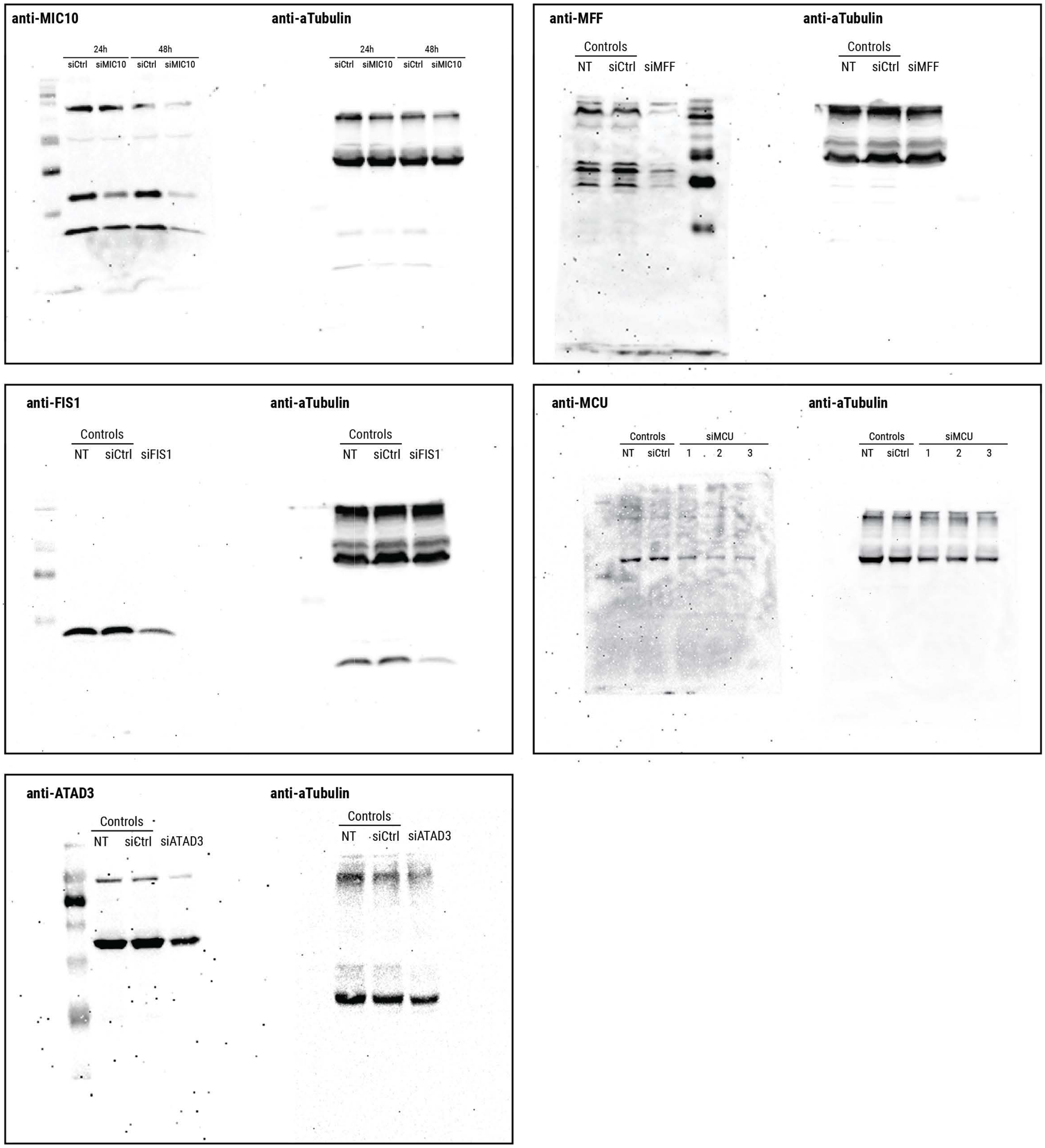


**Extended Data Fig. 3.**

Uncropped western blots depicted in Extended Data Fig. 1-2, along with their loading controls.
